# The limits of scaling in aggregation-driven patterning of cell collectives

**DOI:** 10.64898/2026.03.27.714601

**Authors:** Michael L. Zhao, Jan Rombouts, Anniek Stokkermans, Anna Erzberger, Alexander Aulehla

## Abstract

Robust development requires maintaining correct proportions as overall size varies. What controls and limits this scaling remains poorly understood due to the complex interplay of mechanical and biochemical embryonic factors. Combining confinement of dissociated embryonic presomitic mesoderm cells with imaging and chemical perturbations, we identified cell aggregation as the initial event in *de novo* posterior pole formation. Using a continuum model based solely on cell-cell attraction, we quantitatively map how the time available for aggregation-driven patterning limits the maximum scalable system size. Small systems allow rapid, robust pattern scaling, whereas coarsening dynamics substantially delay scaled patterns in large systems. Our experiments quantitatively confirm these predicted scaling regimes. Together, our results reveal a developmental time-size tradeoff governing the scaling of aggregation-driven patterns.

## 1 Introduction

How cell fate patterns change with the size of multicellular systems is a fundamental question. Developing embryos exhibit the remarkable ability to scale cell fate patterns with overall system size (1–5). Such scaling enables downstream morphogenetic processes to generate proportionately organized tissues despite variation in embryo size (6–8). This important property of living systems raises a series of interconnected questions. What mechanisms underlie cell fate patterning and determine its ability to scale? How are the size limits to pattern scaling established?

Considerable progress has been made in identifying mechanisms that enable patterns to scale with overall system size (9–11). Morphogen gradients are among the best-studied examples, notably the Bicoid gradient in *Drosophila* embryos (3). These studies have revealed diverse strategies that couple patterning to system size, including source–sink arrangements (12), size-dependent reaction or transport processes (1, 13), feedback regulation (14), and changes in cellular signaling responses (15). Related size-coupling mechanisms can also modify self-organized reaction–diffusion systems, allowing their intrinsic patterning length scales to adjust with system size (16).

Multicellular patterns can also emerge through cell movements, which play important roles across diverse biological contexts (17–25). Cell aggregation is one common form of movement-driven patterning, occurring across both unicellular and multicellular organisms. Examples include aggregative multicellularity in holozoans, chemotactic aggregation in *Dictyostelium*, neutrophil swarming, and precartilage condensation during chondrogenesis (26–29). Because movement-driven patterns depend on cells physically redistributing in space, changes in system size may alter both the distances over which cells must move and the number of cells whose movements must be coordinated, with potential consequences for the spacing and position of the final pattern. Compared with gradient-mediated patterns, it remains less understood how movement-driven patterns become coupled to system size. In particular, it is unclear whether this coupling requires a dedicated size-dependent regulatory mechanism, analogous to expander-like feedback in reaction–diffusion systems, or whether pattern scaling can arise without explicit size-dependent regulation.

Defining the size range over which scaling holds can help distinguish among scaling strategies and reveal the constraints that patterning systems face as tissue size changes. However, experimentally mapping both the scaling regime and its limit remains challenging. Studies of naturally varying and experimentally manipulated embryos have demonstrated robust scaling across embryo sizes that remain compatible with development (1, 2, 8). These studies reveal how scaling operates within that range, but provide more limited access to the broader size variation needed to determine where scaling breaks down. Tissue regeneration provides one route to study pattern scaling over larger size ranges than those observed during normal development (30). *In vitro* models derived from primary tissue or cultured embryonic stem cells provide a complementary approach by enabling experimental manipulation of size, density, and spatial confinement in ways that are difficult to achieve in native biological contexts (31–40). These systems therefore enable system size to be varied systematically while patterning outcomes are measured, allowing scaling and its limits to be tested in a controlled setting.

In this study, we adapted an *ex vivo* model of embryonic posterior pole formation (35, 41) to investigate how patterns change with system size. Imaging and chemical perturbation experiments identified cell aggregation as a key early process during pattern formation. We used this observation to formulate a continuum description of cell-cell attraction, which predicts specific time-dependent limits of pattern scaling. These findings quantitatively resolve how the finite time available for aggregation-driven patterning limits the system size over which pattern scaling can be achieved, revealing a tradeoff between the timescale of cell-attraction-based patterning and the spatial range over which scaling remains accessible.

## 2 Results

### 2.1 Limits to scaling are observed in *ex vivo* posterior fate patterns

Dissociated cells derived from mouse embryonic presomitic mesoderm (PSM) form *de novo* structures termed emergent PSMs (ePSMs) (Fig. 1A, 35, 41). ePSMs have been previously shown to recapitulate key features of posterior fate organization in the PSM, including expression of the posterior marker Brachyury (*T*) in a radial arrangement in which central positions correspond to more posterior identities and peripheral positions to more anterior identities (35). We exploited the approximately two-dimensional organization of ePSMs to establish an imaging workflow that captured both single-cell behavior and the mesoscale *T* expression pattern. Combined with micropatterning to control cell attachment area, this system provided a tractable setting in which to test how posterior fate patterns depend on system size.

**Figure 1:**
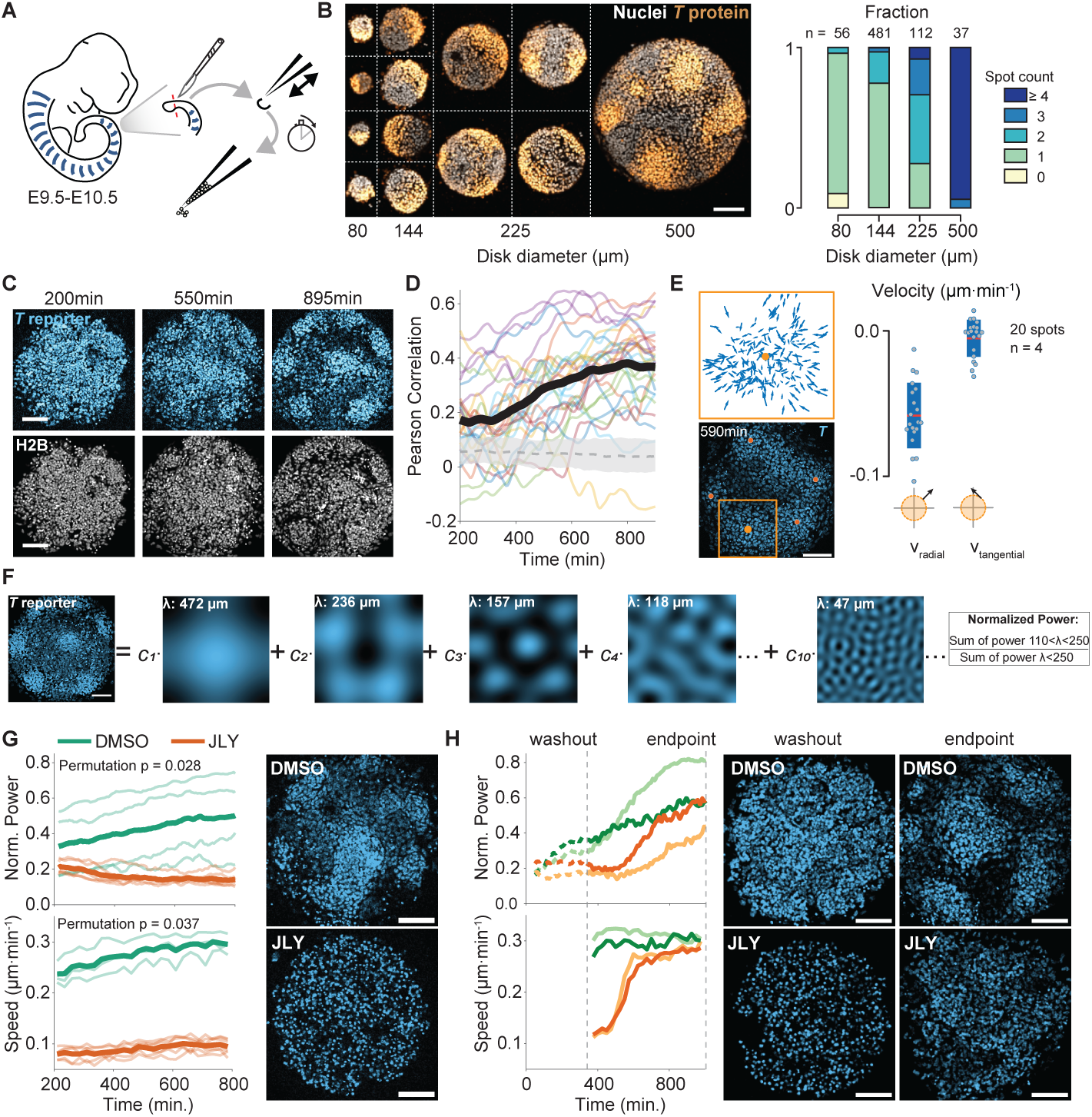
Limited scaling of *T* patterns in the ePSM assay. **A)** We dissociate PSM tissue from mouse embryos as schematically shown, and define time 0 min in all subsequent analyses as the point of dissociation. **B)** We visualized *T* protein levels by immunofluorescence staining (nuclei: DAPI, *T* protein: Alexa-Fluor 568) in PSM cells on disks of different diameters at *t* = 1, 140 min post dissociation (left: example composite image formed by manual tiling, right: quantification of *T* spot numbers). Scale bar = 100 µm. **C)** Example montage shows the dynamic progression of *T* patterning after dissociation of PSM cells. Scale bar = 100 µm. **D)** The Pearson correlation between local *T* activity and local cell density for 27 wells, represented by thin colored lines, increases over time. Mean of all wells is shown by the bold black line. Local areas were obtained by discretizing images onto a 50 µm *×* 50 µm square grid. Gray dashed line and band represent median and 95th percentile bands of a null relationship obtained by random rotations and reflection of the local *T* activity field relative to the local cell density field. **E)** An example of quantified cell velocity vectors obtained from single cell tracking are shown in a 200 µm *×* 200 µm region (orange square) around one of five manually identified *T* spots (orange dots) from 200 to 590 minutes. The time-averaged radial velocities of the cells around *T* spot centers collected from four different wells (right) are shown together with the mean (red line) and standard deviation (blue box). **F)** To quantify *T* patterning dynamics, we compute the normalized power of the *T* reporter signal, defined as shown schematically. Scale bar = 100 µm. **G)** Effect of JLY treatment on the normalized power and cell speed compared to control (DMSO) samples (thin lines: individual well trajectories; bold lines: mean; p-values: permutation test of the treatment *×* time interaction for power and permutation test for difference in average starting speed). Cell speed and normalized power are calculated for the same samples with the exception of 1 day that did not have matched cell tracking data. Images represent examples for each treatment condition, scale bar = 100 µm. **H)** Effect of JLY washout on the normalized power and cell speed. Images represent examples for each treatment condition before and after JLY washout, scale bar = 100 µm

To determine how patterning depends on system size, we seeded dissociated PSM cells on fibronectin-coated disks with diameters ranging from 80 µm to 500 µm. Cells were fixed after either 4 or 19 hours and stained for *T* protein. Samples fixed after 4 hours confirmed that dissociation generated approximately homogeneous spatial *T* profiles (Fig. S1A). By 19 hours, cells had formed discrete spot-like *T* patterns (Fig. 1B). The number of *T* spots increased with system size: for the larger disks (225 µm and 500 µm), the dominant outcome was a multi-spot pattern, whereas for the smaller disks (80 µm and 144 µm), the dominant outcome was a single *T* spot (Fig. 1B, Fig. S1B). Restricting the analysis to single-spot patterns, we found that spot diameter and area increased with disk size (Fig. S1, B–D). Thus, ePSMs exhibit a limited scaling regime in which a unipolar *T* pattern scales with system size, but above a critical size the dominant outcome shifts to multipolar patterns. The size at which this happens defines the boundary of the scaling regime. This size dependence on simple disk geometries resembles recent observations in other models of A–P patterning, suggesting that shared patterning principles may operate across these systems (42, 43). To understand the basis of this scaling limit, we next investigated the dynamical mechanisms underlying *T* pattern formation.

We performed time-lapse fluorescence imaging of dissociated PSM cells from a *T* reporter mouse line to capture the onset of pattern formation (Movie S1, (44)). Western blotting and single-cell analysis confirmed that the reporter faithfully reflected *T* protein levels throughout the patterning time course (Fig. S2A–C). We imaged cells on fibronectin-coated disks of approximately 450 µm diameter, a system size well above the scaling limit and within the multi-spot regime. Following a 200-minute period to allow for cell attachment and spreading, we observed dissociated cells forming spot-like *T* patterns that were accompanied by the emergence of dense cell clusters (Fig. 1C, Fig. S3A,B). The average nearest-neighbor distance between manually identified *T* spots was 164 µm and remained consistent across the wells observed, in agreement with previous ePSM measurements (Fig. S3C,D, 35). Over the course of pattern formation, local *T* reporter activity became positively correlated with local cell density (Fig. 1D). The appearance of dense clusters, together with this positive correlation, suggested that cell movements contribute to *T* pattern formation. To examine this possibility, we tracked cell nuclei during the first half of the experiment, between 200 and 590 minutes post-dissociation. To determine how cells incorporated into developing *T* spots reached these regions, we identified cells located within 60 µm of manually identified spot centers at the end of the tracking interval and reconstructed their preceding movements. These cells showed a pronounced inward radial bias, consistent with cellular aggregation (Fig. 1E). Consistent with this interpretation, particle image velocimetry showed that cell flows across the well tended to converge on future sites of high *T* gene activity (Fig. S4A–D). To determine whether *T* levels at the beginning of the tracking interval were associated with aggregation propensity, we asked whether *T* reporter activity at 200 minutes post-dissociation predicted subsequent aggregation-directed movement. Cells with above-median initial *T* reporter activity showed stronger inward radial movement toward future spot centers than cells with below-median reporter activity, whereas their tangential velocities did not differ (Fig. S4E,F). Directed, aggregation-like cell movements therefore accompany the initial formation of *T* patterns and are stronger among cells with higher initial *T* reporter activity.

To test whether cell movements are required for the initial formation of regions marked by *T*, we applied a pharmacological cocktail chosen to broadly suppress movement within the cell population. Specifically, we used the inhibitor combination Jasplakinolide, Latrunculin A, and Y-27632 (JLY), previously described to arrest actin dynamics (45). To quantify the effect on spatial patterning, we used Fourier analysis to decompose *T* reporter images into spatial modes, each associated with a characteristic wavelength (Fig. 1F). We then defined a normalized power score as the summed contribution of modes with wavelengths between 110 and 250 µm, consistent with the observed nearest-neighbor spacing, divided by the total power across all modes of 250 µm or less (Fig. 1F). JLY treatment strongly reduced both cell speed and pattern progression, as quantified by cell speed and normalized power, respectively (Fig. 1G). Washout after 6 hours of JLY treatment showed that both effects were partially reversible following media replacement (Fig. 1H, Movies S2,S3). Notably, cell speed began to recover before *T* patterning resumed, consistent with cell movement acting upstream of visible *T* pattern formation. These experiments provide evidence that aggregation-like cell movements are required for the initiation of *T* pattern formation.

### 2.2 Cell aggregation precedes changes in *T* expression

Although the previous experiments identified a role for aggregation in *T* patterning, it remained unclear whether changes in *T* expression due to additional signaling-dependent effects occurred concurrently or instead followed a specific temporal order. As a readout of signaling state, we therefore quantified cell-level *T* expression during ePSM pattern formation. Over the course of the experiment, the distribution of *T* expression across the cell population broadened while the mean *T* level changed comparatively little. This indicates that some cells increased whereas others decreased *T* expression over time (Fig. 2A). This widening of the expression distribution became most apparent in the second half of the experiment, beginning around 500–600 minutes post-dissociation.

**Figure 2:**
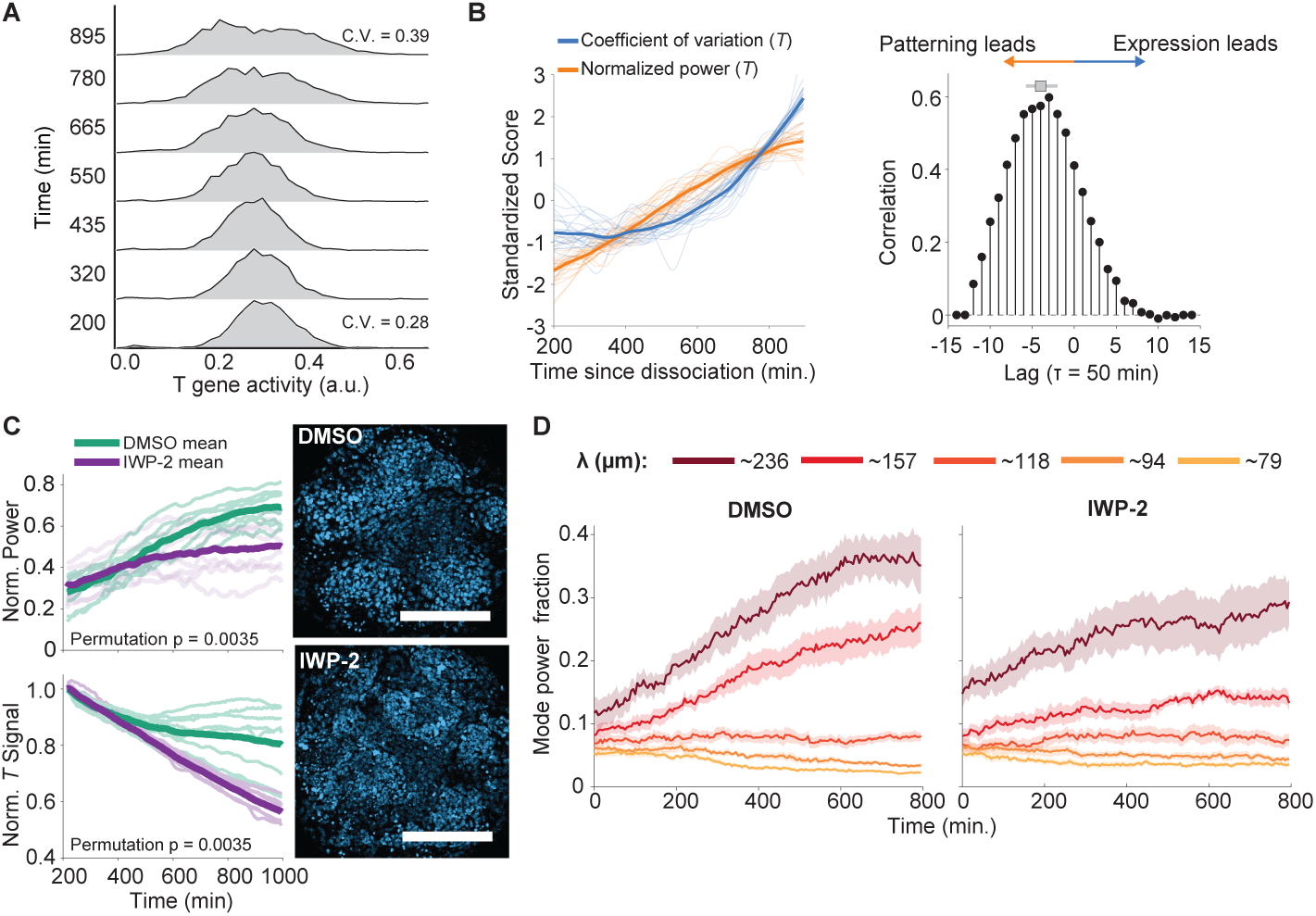
Cell aggregation precedes changes in *T* gene activity. **A)** Time evolution of the *T* gene activity distribution across the population of cells in one well. **B)** Comparison of standardized scores of the normalized power of cell nuclei patterns quantified from *T* reporter images (orange) to the *T* distribution coefficient of variation (blue) for 27 wells. Cross-correlation analysis, averaged across wells, exhibited a peak lag at *−*150 minutes. (Gray square and line: median and interquartile range of cross-correlation peak positions of the individual wells.) **C)** Effect of IWP-2 treatment on the normalized power and normalized *T* signal compared to control (DMSO) samples (thin lines: individual well trajectories; bold lines: mean; p-values: permutation test of the treatment *×* time interaction for both normalized power and normalized *T* signal). Images represent examples for each treatment condition and do not share the same intensity scale, scale bar = 200 µm. **D)** Time courses of spatial mode fractions in DMSO and IWP-2 conditions following Fourier decomposition. Bold lines represent mean of *n* = 8 replicates per condition and shaded regions are S.E.

We next compared these cell-level expression changes with the onset of spatial pattern formation by performing a cross-correlation analysis between the coefficient of variation (CV) of single-cell *T* expression and the normalized power of the emerging pattern. Relative to normalized power, the increase in *T* CV was delayed by several hundred minutes (Fig. 2B, Fig. S5A), indicating that the first detectable evidence of pattern formation appeared before substantial divergence in cellular *T* expression. We then repeated the analysis using normalized power quantified from images of the constitutively expressed H2B-mCherry to compare expression changes more directly with density changes resulting from the aggregation-driven component of patterning (Fig. S5B and S6A). Again, the increase in normalized power preceded the increase in *T* CV by several hundred minutes.

We additionally asked whether the delay in CV could be the result of a reporter-specific delay. We repeated a similar analysis using dissociated cells from an endogenous knock-in reporter line for *β*-catenin, a key WNT-signaling effector and upstream regulator of *T* expression (46). Nuclear *β*-catenin, indicative of active WNT signaling, changed its spatial pattern and cell-to-cell variation on timescales similar to those observed for the *T* reporter (Fig. S6B–E). Together with the strong correspondence between *T* reporter and *T* protein levels throughout patterning (Fig. S2C), these results argue that the delayed increase in cell-to-cell *T* variation is not readily explained by an intrinsic delay in the *T* reporter.

Finally, we asked whether the late increase in *T* expression observed in a subset of cells is required for pattern formation by blocking WNT ligand secretion throughout the experiment with IWP-2 (47). IWP-2 reduced the mean *T* reporter signal but did not prevent the emergence of spatial patterning (Fig. 2C). However, patterns in IWP-2-treated wells appeared more fragmented than in controls (Fig. 2C, Fig. S7A,B). This disruption was corroborated by immunostaining for *T* protein, which further suggested that the correlation between cell density and *T* expression was weakened under these conditions (Fig. S7C,D). Consistent with this fragmented appearance, analysis of individual Fourier modes showed that IWP-2 flattened the late rise of the largest spatial modes associated with the characteristic *T* spot spacing (Fig. 2D). Together, these results suggest that WNT-signaling-dependent increases in *T* expression act to reinforce, rather than drive, the developing pattern.

Collectively, our dissection of *T* patterning dynamics supports a temporal sequence in which aggregation drives early spatial patterning, whereas subsequent WNT-signaling dependent changes reinforce the developing *T* pattern. This temporal ordering is qualitatively consistent with recent observations in gastruloid models of A–P patterning (48, 49), indicating that similar dynamical ordering can arise across distinct *in vitro* models of A–P patterning.

### 2.3 Quantifying the biophysical parameters of *T* cell aggregation using nonlocal continuum modeling

Given the central role of aggregation in *T* patterning, we sought to theoretically determine if this mechanism alone accounts for our observations. Aggregation-driven pattern formation requires that directed cell motion due to long-range cell-cell attraction overcomes undirected, random cell movements (20, 50, 51). To quantitatively characterize the random and directed motion of *T*-expressing cells in our system, we analyzed the early dynamics of *T* patterning using a nonlocal continuum theory (52–54). Motivated by the observed relationships between *T* reporter activity, local cell density, and aggregation-directed radial movement (Fig. 1D and Fig. S4E,F), we use a minimal model describing the dynamics of the high-*T*-expressing population, herein referred to as *T* +. The equation describing *ρ*, the density of *T* + cells, reads

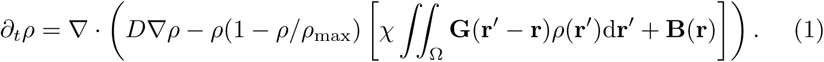

In this framework, an effective diffusion coefficient *D* sets the strength of random cell movements, and cell-cell attraction between *T* + cells is described by a nonlocal term, parameterized by an interaction strength *χ*. The interaction kernel **G** describes how cell-cell interaction strength depends on distance. In this work, we use an exponentially decaying interaction with characteristic range *σ* (Fig. 3A, Supplementary Material). The term 1 *− ρ/ρ*_max_ is called a packing term, which ensures that local velocity goes to zero when the density approaches *ρ*_max_ and prevents aggregation to very high densities (55). The cells move within a confined space denoted by Ω, and the function **B** describes interaction effects with material outside of this space. Here, we use so-called neutral boundary effects, which assume the outside material has an attractive effect similar to that of the cells (see Supplementary Theory).

**Figure 3:**
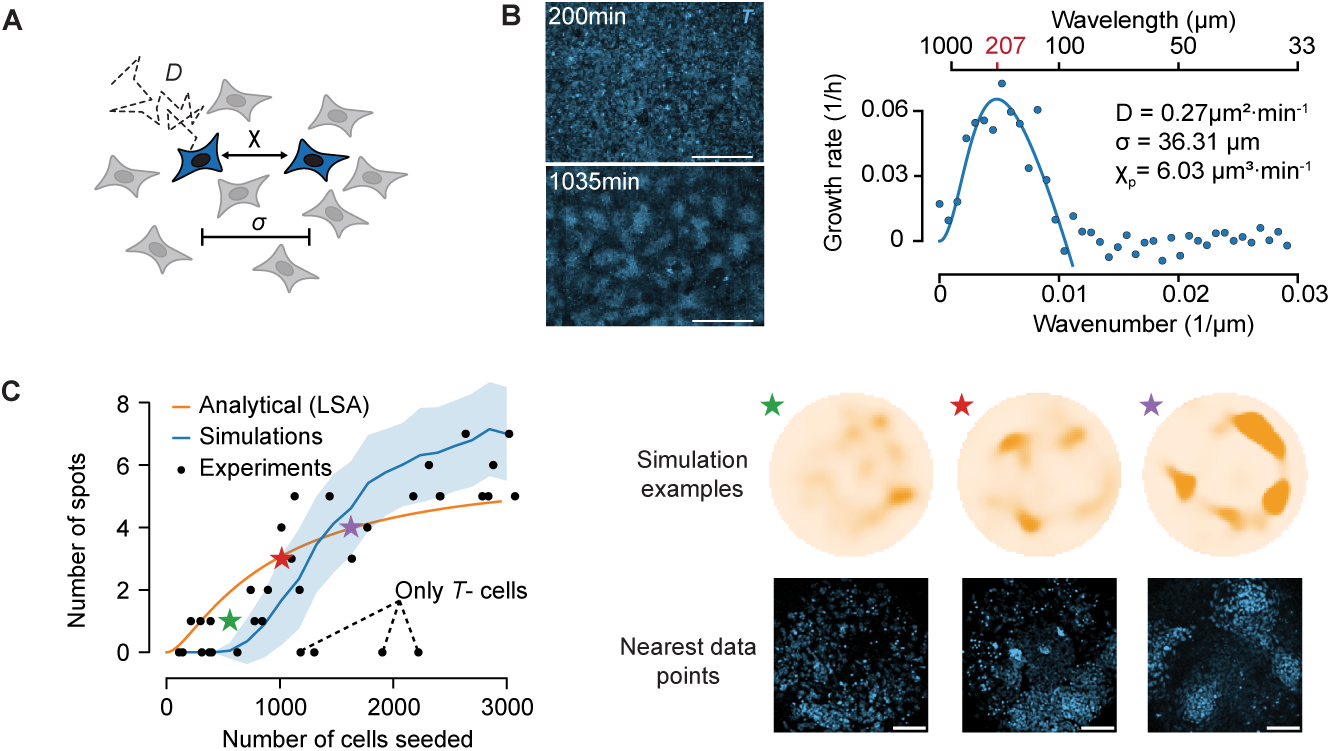
Cell movements determine *T* patterning dynamics. **A)** The theory considers the distribution of *T* + cells (blue), which move with a random component (effective diffusion coefficient *D*) and a directed component due to cell-cell attraction with strength *χ* and range *σ*. **B)** Fitting the experimental dispersion relation from patterning experiments performed on 2 mm *×* 1.5 mm wells (left panels) allows us to estimate the cell movement parameters (best-fit values for one out of three replicates shown). Scale bar = 500 µm. The wavelength indicated in red is the one with maximal growth rate according to the fitted curve. **C)** Simulations on 500 µm wells with different initial densities show spot counts similar to experiments with varying cell seeding number and initial *T* + cell fraction. Shaded area: standard error of simulations. Stars mark selected example simulations, shown together with the closest experimental images. Three wells contained only anterior cell fates with low *T* activity. Scale bar = 100 µm. An analytical estimate based on the linear stability analysis provides an estimate for the number of spots to expect in the absence of nonlinear effects (orange line).

At the onset of patterning, when the density distribution is close to a spatially uniform profile, a linear stability analysis can be used to investigate the dynamics and typical length scales of emerging patterns. In particular, the spatial distribution of *T* + cells (Fig. S8) can be decomposed into spatial modes, each with a specific wavelength — similar to the analysis used to quantify patterns in Fig. 1F. The linear stability analysis allows one to calculate which of these modes grow or decay, and thus are expected to contribute to the pattern. Calculating the growth rate as a function of the mode’s wavelength yields a dispersion relation, which itself depends on the biophysical parameters of cell movement (Supplementary Theory, Fig. S9A).

To fit this theoretical relation to experimental measurements, we observed patterning dynamics in larger wells (2 mm x 1.5 mm) where boundary effects are negligible (Movie S4). Fourier analysis of the *T* + cell density field over time showed that the spatial modes corresponding to patterning grew exponentially for a 500 minute interval (Fig. S9B–F). This allowed a growth rate to be fitted to each mode and the experimental dispersion relation for each replicate well to be determined.

Fitting the theoretical curve to the experimentally obtained dispersion relation, we inferred the biophysical parameters underlying aggregation-driven patterning independently for each well. Across *n* = 3 experiments, we obtained an average effective diffusion coefficient of *D* = 0.25 *±* 0.07 µm^2^ min*^−^*^1^, an average cell-cell attraction strength of *χ* = 5.8 *±* 0.9 µm^3^*/*min*/*cells, and an average interaction range of *σ* = 41 *±* 8 µm (Fig. 3B, Fig. S9G,H, Fig. S10).

The magnitude of the random component, *D*, is similar to values obtained for cell movement in crowded environments for other multicellular systems (56–59). The units of *χρ*_0_ are those of a velocity, and the maximal value of the speed at a point in our model due to an attracting region of constant density *ρ* is 2*χρ* (Supplemental Theory, Fig. S11A–C). We thus expect *χρ*_0_ to give an order-of-magnitude estimate of typical cell velocities. Using the estimated *ρ*_0_ from experiments and the fitted value of *χ*, the product of the two values is 0.04 µm min*^−^*^1^. This estimate of the velocity component due to cell-cell interactions is consistent with radial velocities obtained from single cell tracking (Fig. 1E). At the fitted interaction range, *σ*, the interaction strength has decayed to approximately 37% of its maximum at 4.8 *±* 1.0 nuclei spacings and 10% of its maximum at 11 *±* 2 nuclei spacings, indicating that cell-cell interactions extend beyond immediate neighbors (Fig. S11D–F). The fitted parameter values place the system well within the pattern-forming regime predicted by the theory (Fig. S12A). The fastest growing modes of the fitted dispersion relations correspond to pattern wavelengths of 220 *±* 30 µm, differing from the mean value of the nearest-neighbor distance between *T* spots by 1.8 standard deviations (Fig. S9F–H, Fig. S3C). A scaling analysis shows that the initial pattern’s wavelength is, to leading order, proportional to the interaction length *σ*. Moreover, the pattern’s growth rate is inversely proportional to *σ* (Fig. S12B–C, Supplemental Theory). This result, applicable to large systems, agrees with the intuition that to establish large-wavelength patterns, cells need to move across larger distances, which takes more time.

To validate our parameter estimates, we performed numerical simulations of the continuum equation using the fitted values without additional free parameters for different initial cell densities, and compared the results to experiments in which we varied the *T* + fraction and total cell seeding number (Fig. S13A–C). We obtained a close quantitative agreement between simulations and data, confirming the predictive power of the inferred parameters (Fig. 3C). Moreover, these results qualitatively agree with the analytically obtained result that increasing the density makes the wavelength of the pattern smaller, which means more spots on a given system size (Fig. S12D, Supplemental Theory).

In summary, both analytical and numerical results support our experimental findings that aggregation-driven patterning underlies the initial distribution dynamics of *T* + cells. Our estimates of motility and interaction parameters show that this system is robustly capable of spontaneously forming density patterns.

### 2.4 Aggregation dynamics couple system size to patterning timescales

While quantifying the patterning dynamics in large wells allowed us to characterize the pattern-forming mechanism in the absence of boundary effects, the size and shape of the space within which patterning takes place can strongly influence its outcomes (60). The effects of system size, shape, growth and deformations have been extensively studied for (mechano)-chemical patterning systems (61–65). For example, a generic effect of size is that, in confined spaces, only a *discrete set* of pattern modes whose wavelengths fit within the available space can develop (66, Chapter 2). With estimates of all model parameters in hand, we used simulations to investigate pattern formation on small disks, and generated fit-free predictions of patterning outcomes as a function of system size to compare with the results of Fig. 1B. We found multi-spot patterns on larger disks, and unipolar patterns on a range of smaller disks (Fig. 4A). Using initial conditions based on experimental density profiles at 200 minutes, the independently parameterized simulations showed good agreement with experiments in the spot-count distributions for 144, 225, and 500 µm disks with the main discrepancy occurring for the 80 µm disks (Fig. 4B). In particular, experiments showed a scaling regime between 80–144 µm, while in our simulations the scaling regime — defined as at least 50% of simulations showing a unipolar outcome — was 100–160 µm. In simulations with a unipolar outcome, the spot size roughly scales with the system size similarly to the experimental result (Fig. S14).

**Figure 4:**
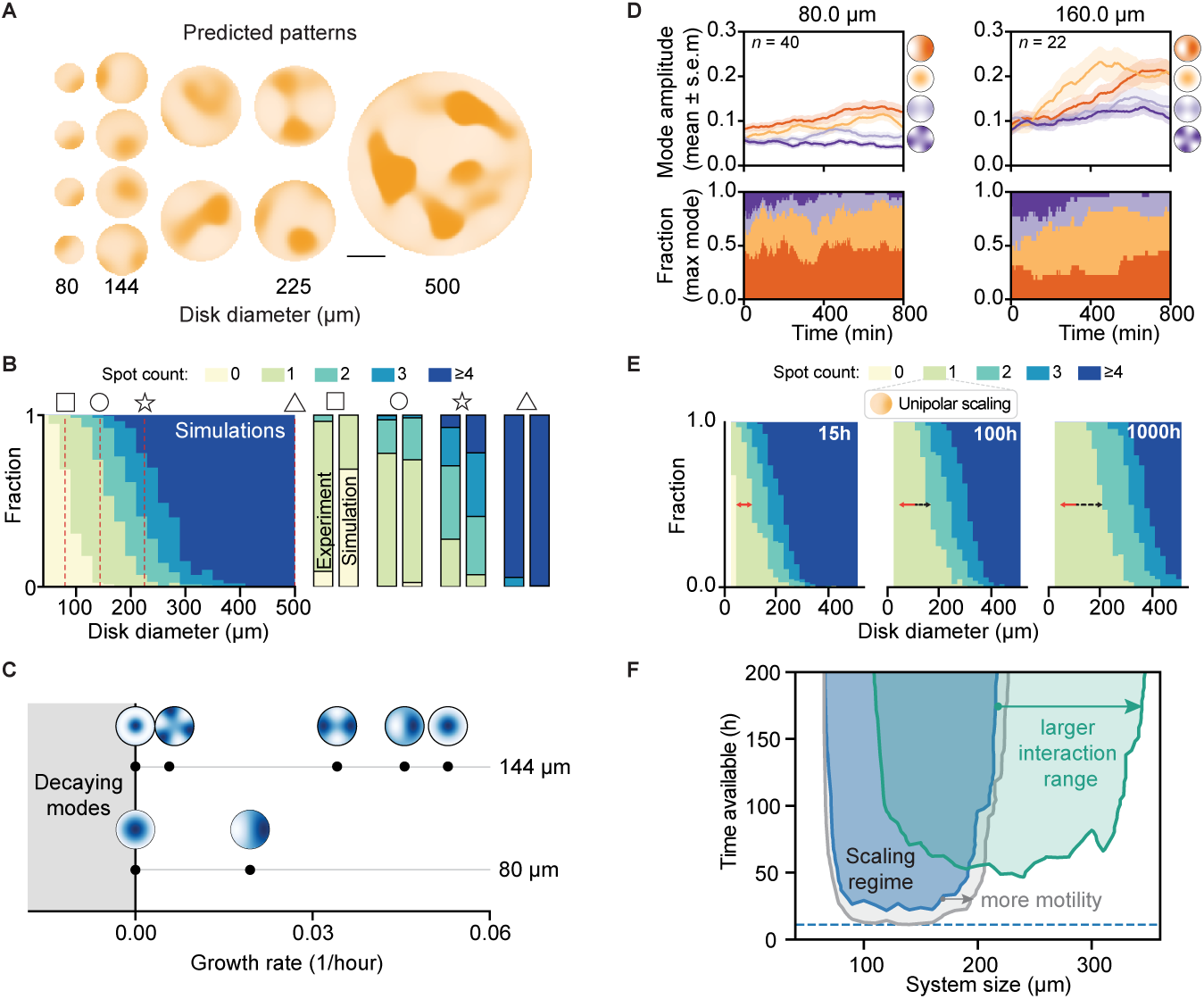
Finite-size and non-linear effects explain patterning outcomes across system sizes. **A)** Representative simulations show patterning outcomes for disks of different sizes at *t* = 1, 100 min. Scale bar = 100 µm. **B)** The distribution of spot numbers across simulations changes as a function of disk size. For the experimentally realized disk sizes (red dashed lines and icons), we compare spot count statistics between simulations and data shown in Fig. 1B. **C)** Growth rates and shape of spatial modes with zero or positive growth rates obtained from a linear stability analysis on disks. **D**) We project the time-lapse disk data onto four selected modes and compute their amplitude (top, cross-disk average) and the distribution of the dominating modes (bottom) over time. Shaded area is S.E.M. **E)** Spot count statistics for simulations conducted past typical experimental durations show how the scaling region of the unipolar pattern increases (black dashed arrow, with red arrow referencing the scaling width at 15 hours). **F)** Performing simulations across time windows and system sizes (100 runs per point), we compute the unipolar scaling regime for the experimental ePSM parameters (blue region), for a two-fold increase in cell motility *D* and *χ* (gray region), and a two-fold increase in the interaction range *σ* (green region). We define the scaling region as the set of points for which at least 50% of simulations exhibit a single spot.W

To identify the mechanisms underlying this scaling behavior, we performed a linear stability analysis adapted to circular domains (Supplemental Materials). In contrast to infinite domains, where Fourier modes are appropriate, in this case the modes change with domain size and parameter values. Our analysis showed that initial patterning dynamics can be described by only a few growing spatial modes (Fig. 4C). For 80 µm disks, only a single spatial mode can grow: the unipolar pattern. For 144 µm disks however, single-spot and two-spot patterns have similar growth rates, indicating that initially multiple spots may form. The size of the system thus strongly constrains the possible patterns.

To investigate whether the temporal dynamics of the experimental system can be described by these unstable modes, we collected a dataset of time-resolved *T* patterning dynamics on small disks. After filtering low density disks and disks with cells that enter at later time points, we decomposed the remaining observed density patterns into spatial modes. We used the modes obtained by the linear analysis and analyzed the contributions of four modes: central spot, unipolar, bipolar, and tripolar (Fig. 4C,D; Fig. S15A,B; Movie S5). These results confirm a number of predictions from the linear stability analysis. First, for 80 µm diameter disks, the polar mode is dominant and growing, but its growth rate is smaller than the dominant growing modes on the larger disks. Additionally, for 160 µm diameter disks, both central-spot and polar modes have large growth rates, whereas two- and three-spot modes grow at a slower rate, also consistent with the linear analysis.

At later times, the polar mode becomes dominant on the 160 µm disks and the other modes drop in amplitude, suggestive of nonlinear effects. Indeed, on some 160 µm disks a cell cluster that formed in the center drifted to the side at later times (second example in Fig. S15B). This observation is consistent with our previous theoretical investigations of one-dimensional aggregating systems that show the same behavior (54).

In larger systems imaged over longer periods, we also directly observed the joining of initially separate *T* + spots (Movie S6). Such merger events are characteristic of coarsening, a non-linear phenomenon, in aggregation-driven systems (67, 68). Coarsening can in principle enable the scaling of a unipolar pattern, i.e. leading to a single *T* + domain, across arbitrarily large systems. However, the time required to reach this state increases nonlinearly with system size because the intermediate multi-spot patterns are long-lived transients in the system dynamics (54).

To investigate quantitatively how patterning duration influences this scaling capacity for aggregating multicellular systems, we investigated spot-count statistics as a function of system size in long-term simulations of up to 1,000 h. For such long simulations, we observed an approximately 2.5-fold increase in the width of the unipolar scaling regime, confirming that the time available for a given patterning process effectively acts to limit the size range over which patterns can scale (Fig. 4E). Conducting simulations over a wide range of system sizes and developmentally relevant time windows, we mapped out the full unipolar scaling regime for the aggregation parameters of the experimental system (Fig. 4F, blue region), and found that predominantly two effects determine the shape of the scaling regime. First, the patterning timescale imposed by the parameters *D, σ* and *χ* (horizontal dotted line) sets a lower bound for patterning time. Second, the finite size effects and nonlinear coarsening effects shape the boundaries of the scaling regime on longer timescales. To see how the scaling regime depends on the biophysical parameters controlling aggregation dynamics, we simulated the effect of a two-fold increase in the motility parameters *χ* and *D*, and the interaction range *σ*. We found that increasing motility expanded the scaling regime modestly in all directions, mainly due to the global speedup of all cell motion, which mathematically corresponds to a rescaling of time. A larger interaction range, on the other hand, increased the time required to develop a pattern while shifting and expanding the size range over which unipolar patterns dominate due to a combination of finite-size and long-term nonlinear effects. Since the dominant growth rate is inversely proportional to *σ*, it is indeed expected that systems with larger interaction ranges develop patterns more slowly (Fig. S12C).

Together, our results demonstrate how aggregation dynamics links patterning time scales to system size and delineates mechanisms for controlling the scaling of unipolar patterns.

## 3 Discussion

Primary cells derived from the PSM retain a robust ability to self-organize into structures characterized by multipolar *T*-positive domains, a pattern that departs from the unipolar *T* domain observed *in situ* (Fig. 1B, 35). We harnessed this potential to address how cell fate patterns depend on system size and identified a regime in which unipolar *T* patterns reliably form and scale to overall size. Experiments and theory revealed a key principle setting the size–patterning relationship in aggregating systems: scaling is robust for small system sizes, but for larger systems the time required to achieve scaling exceeds realistic developmental timescales. The underlying attractive cell-cell interactions therefore impose a time–size constraint on pattern scaling, setting an upper system size that can achieve a unipolar outcome within a limited patterning window. Moreover, because the theory depends on an effective interaction rather than on a specific mechanism, the resulting constraint applies to any mechanism that produces comparable aggregation dynamics.

Importantly, this scaling limit is not fixed but reflects a fundamental time–size tradeoff. For a given developmental time window, only a restricted range of system sizes successfully achieves unipolar scaling. Both linear and nonlinear effects arising from aggregation-driven patterning shape this tradeoff: finite-size effects limit the possible number of initially forming aggregates, delineating a range of system sizes in which unipolar patterns are the only ones that can form. Beyond this range, larger systems support initial patterns with multiple aggregates that subsequently undergo slow merging toward a final unipolar state — an outcome that may not be reached within the available time window. Our results therefore suggest that system size must be constrained in accordance with the time available to achieve unipolar outcomes.

The size range over which patterns scale can be enlarged by modifying the effective parameters governing cell movement and attraction. Our simulations show that increasing motility primarily accelerates the dynamics, whereas changing the interaction range alters both the characteristic spatial scale and the time required for pattern formation. These parameters could arise from different molecular or cellular mechanisms. In some cases, more detailed descriptions can be mathematically recast as nonlocal advection equations such as the one used here (54, 69). The Keller–Segel model, originally motivated by chemotaxis-driven aggregation in *Dictyostelium*, provides one such example (50) while alternative interaction rules can be represented by different interaction kernels (70). The value of the interaction range *σ* we inferred extends over several cell diameters or neighbor distances and is compatible with longer-range processes such as traveling chemical waves (27, 28), cellular protrusions (21, 71), or stress propagation through the tissue (23, 25). At the same time, local adhesion-mediated sorting may contribute alongside longer-range interactions, as reported in recent studies (22, 48, 49). Perturbing these candidate mechanisms and measuring their effects on random motion, attraction strength, and interaction range would reveal how molecular and cellular regulation shifts the scaling regime predicted by the model.

Whether embryos face comparable time–size constraints remains unknown, but early mouse gastrulation provides a plausible context in which to investigate this possibility *in situ*. Between E6.0 and E6.5, the embryonic portion of the epiblast has linear dimensions of approximately 150 µm to 200 µm, close to the boundary of the scaling regime identified here (72). Across these early developmental stages, Rac1-dependent migration contributes to polarization of the anterior visceral endoderm and directional redistribution of nascent mesoderm, while reported mesodermal cell velocities of 0.1 µm min*^−^*^1^ to 0.3 µm min*^−^*^1^ and approximately 0.7 µm min*^−^*^1^ are similar in magnitude to those measured in our *ex vivo* system (73–75). These similarities in spatial scale and movement rate motivate the hypothesis that movement-driven patterning in the embryo may encounter related time–size constraints.

Within complex tissues such as the developing embryo, aggregation-driven patterning occurs concurrently with cell signaling, proliferation, and global shape changes. How these factors influence pattern scaling depends on the relative timescales between these processes. In the simplified context we studied, chemical inhibitor experiments showed that WNT-dependent regulation altered both *T* expression and spatial patterning, demonstrating that cell-state regulation can modify aggregation-driven patterns. A direct extension of the current model could therefore allow the average *T* + density to change through transitions between high- and low-*T* states, while addition of a cell-turnover term could additionally account for changes in population size. An alternative approach would be to couple spatial patterning to an intracellular gene-regulatory network as was recently done within a reaction-diffusion framework (76). Such extensions would test how regulation of the *T* + population modifies the aggregation-driven time–size tradeoff, including whether changes in cell-state composition stabilize multipolar outcomes rather than permitting eventual convergence to a single domain (77). More broadly, the strong geometric constraints identified in this work raise the question of how living systems use size and shape to steer self-organized patterns. Embryo geometry influences how gastrulation proceeds during development (78), while confinement of organoid cultures is recognized as an important factor for improving reproducibility (79). These examples provide empirical evidence for a guiding role of geometrical constraints and suggest future avenues for harnessing the principles identified here.

## Acknowledgments

We thank Hanspeter Herzel, Jordi Garcia-Ojalvo, David Oriola, Laeschkir Würthner and Takashi Hiiragi for manuscript feedback. We additionally thank James E. Ferrell for advice and discussions throughout the history of this project. This work was supported by Laboratory Animal Resources at the EMBL. We acknowledge the use of the EMBL High Performance Computing resources and we thank the Advanced Light Microscopy Facility (ALMF) at the EMBL and Zeiss for support. MLZ was supported by a Bridging Excellence Fellowship from the EMBL—Stanford Life Science Alliance and a ‘Figure 1 Theory’ Grant from Theory@EMBL Transversal Theme. JR was supported by an EMBL Interdisciplinary Postdoctoral Fellowship (EIPOD4) program under Marie Sklodowska-Curie Actions Cofund (grant agreement no. 847543) and a postdoctoral fellowship from FNRS (Charǵe de recherches). This work was supported by the EMBL and received funding from the European Research Council under an ERC consolidator grant agreement n.866537 to AA. Code and example data available at: https://github.com/JanRombouts/zhao-rombouts-scaling.

## Supplementary Material

Materials and Methods

Supplemental Theory

Figs. S1 to S15

Movies S1-S6

Supplemental References

## Notes

### Competing Interest Statement

The authors have declared no competing interest.

### Summary of Updates

The manuscript has been substantially revised throughout. This version includes a revised introduction to focus on movement driven patterning, additional analysis relating initial T reporter activity to aggregation-directed movement, direct observations of aggregate merging events, expanded presentation and validation of the theoretical model, further analysis of pattern scaling across system sizes, and an expanded discussion regarding biological scope and potential in vivo relevance.

