## Supplementary Material for "The limits of scaling in aggregation-driven patterning of cell collectives"

### Supporting Information

#### 1 Supplemental Materials and Methods

##### Animal Care and Use

Mice used in all breeding and experiments were housed under a veterinarian's supervision in the animal facility at EMBL following the guidelines of the European Commission, revised directive 2010/63/EU and AVMA guidelines 2007. Approval for animal experiments was obtained from the EMBL Institutional Animal Care and Use Committee (project codes: 21–001_HD_AA and 25-014_HD_AA). The detection of a vaginal plug was designated as embryonic day (E) 0.5, and all experiments were conducted with embryos between the ages of E9.5 and E10.5. Pregnant females were euthanized by cervical dislocation in accordance with the approved institutional protocol. Embryos were immediately removed from the uterus and transferred to dissection medium for tissue collection.

##### Mouse Lines and Genotyping

Three mouse lines were used in this study. The first mouse line expressed a Histone-2B-eGFP based Brachyury gene activity reporter, $T^{nEGFP\text{-}CreERT2/+}$, was previously described in (1). The second mouse line expressing a red fluorescent Histone-2B fusion protein from the *Rosa* locus, R26-H2B-mCherry, was previously described in (2). Both fluorescent reporters were maintained together in a single CD-1 strain and all embryos used for experiments were heterozygous for the two transgenic alleles. The third mouse line harbored an endogenous 5' knock-in of the beta-catenin gene with a Venus yellow fluorescent protein. The generation of this mouse used embryonic stem cells from (3) and will be described elsewhere.

R26-H2B-mCherry was genotyped with primers: (5'-TCCCTCGTGATCTGCAACTCCAGTC-3'), (5'-AACCCCAGATGACTACCTATCCTCC-3'), and (5'-GCTGCAGGTCGAGGGACC-3'). Mutant allele band was 270 bp and wild-type allele band was 217 bp.

$T^{nEGFP\text{-}CreERT2/+}$ was genotyped with primers: (5'-GGGCTCTACTTCATCGCATTCC -3'), (5'-TATCCCAGTCTCTGGTCTGTGA-3'), and (5'-TAGGACCCTACCTAGCAAAGGA-3'). Mutant allele band was 400 bp and wild-type allele band was 500 bp.

beta-catenin-venus was genotyped with primers: (5'-GCAATCAGCTGGCCTGGTTT-3'), (5'-ATGGTCCTGCTGGAGTTCGT-3'), and (5'-GCCTAAAACCATTCCCACCC-3'). Mutant allele band was 230 bp and wild-type allele band was 190 bp.

OneTaq 2X Master Mix with Standard Buffer was used (New England Biolabs) for all polymerase chain reactions with protocols according to the manufacturer's protocol. A $T_m$ of 60°C was used for all genotyping reactions.

#### Culture conditions and tissue dissociation

The primary culture medium, referred to as "CM", used during all imaging experiments consisted of DMEM/F12 formulated without glucose, pyruvate, glutamine, or phenol red (Cell Culture Technologies), supplemented with bovine serum albumin (1 % w/v, Equitech-Bio), glutamine ($2\,\text{mmol}\,\text{L}^{-1}$, Sigma-Aldrich), glucose ($2\,\text{mmol}\,\text{L}^{-1}$, Sigma-Aldrich), and penicillin/streptomycin (1 % v/v, Sigma-Aldrich). During embryo dissection and cell dissociation, a modified medium, referred to as "DM", was used, in which antibiotics were omitted, added sodium bicarbonate was reduced from $2.4\,\text{g}\,\text{L}^{-1}$ to $1.2\,\text{g}\,\text{L}^{-1}$, and $17\,\text{mmol}\,\text{L}^{-1}$ HEPES (Gibco) was added.

Cells used in the experiments originated primarily from the tailbud mesoderm region and a small portion of the posterior PSM of mouse embryos. To randomize the cells, a tissue piece spanning the region between the tip of the tail and the beginning of the posterior portion of the presomitic mesoderm was cut away from the embryonic tail using a micro-scalpel. The approximate position of the chordoneural hinge was used as a visual guide for the cut. The dissected tissue was then mechanically dissociated either individually in 1.5 µL of DM or pooled from 4–10 embryos in 4 µL–6 µL of DM before dissociation. Dissociation was performed within one well of a circular 4-well silicone micro-insert (Ibidi) by holding a 10 µL pipette tip (Gilson) against the wall of the well and gently pushing the tissue pieces back and forth through the small opening. The cell mixture was then passed through a 40 µm cell strainer (Falcon) to remove large tissue clumps. The filtered mixture was deposited in micro-wells or on micropatterned surfaces and centrifuged at 400 rcf for 30 s. After centrifugation, DM was aspirated and replaced with 10 µL of CM in each well. After the last well was filled, the entire insert was filled with 150 µL of CM. An additional evaporation barrier was created by filling the coverslip well with water. Time 0 in all experiments is defined at the point of tissue dissociation and cells were then allowed a settling and attachment period for 200 minutes prior to the beginning of analysis. The settling period typically consisted of incubation at 37 °C and 5 % $CO_2$ for 0.5 h–2 h followed by the remaining time in the microscope.

Cells dissociated from individual embryos were seeded into circular micro-wells with a diameter of 400 µm. To prepare the micro-wells, 4-well silicone micro-inserts (Ibidi) were mounted in the center of a one-well chambered #1.5H glass slide (Cellvis) in a vacuum chamber (VWR). For large-well experiments using cells dissociated from multiple embryos, a similar protocol was used, except that the circular micro-wells were replaced by rectangular micro-wells with dimensions of $2\,\text{mm} \times 1.5\,\text{mm}$. The 4-well micro-inserts (Ibidi) remained the same size.

All culture surfaces were coated with fibronectin using one of two protocols. In the first protocol, used for micro-wells, a small drop of fibronectin solution ($0.2\,\text{mg}\,\text{mL}^{-1}$) was deposited on top of the culture surface and incubated for 10 min at 37 °C. After incubation, the fibronectin solution was removed and followed by 2–3 washes with 10 µL of DM. In the second protocol, used for micropatterned surfaces, 10 µL of a $0.05\,\text{mg}\,\text{mL}^{-1}$ fibronectin solution was deposited on top of the surface and incubated for 2 h at 37 °C, followed by 2–3 washes with 10 µL of DM. In all cases, special care was taken not to touch the imaging surface after fibronectin coating.

#### Micropatterning - Fixed Cell Experiments

Glass coverslips ($20\,\text{mm} \times 20\,\text{mm}$) were cleaned by placing them in a rack, immersing them in 70 % ethanol, and sonicating for 10 min at 4 °C, followed by air-drying overnight in a laminar-flow hood. Coverslips were then plasma-treated (∼1 min) and immediately passivated by placing the activated side onto 100 µL drops of PLL(20)-g[3.5]-PEG(2) (PLL-PEG, SuSoS) prepared at $0.1\,\text{mg}\,\text{mL}^{-1}$ by a 20-fold dilution of a $2\,\text{mg}\,\text{mL}^{-1}$ stock in $10\,\text{mmol}\,\text{L}^{-1}$ HEPES buffer (pH 7.4) on Parafilm, and incubated for 1 h at room temperature. After incubation, coverslips were rinsed by dipping in ultrapure water and dried.

Micropatterns were generated by UV–ozone degradation of the PLL-PEG layer using a chrome photomask in a UV–ozone oven. Briefly, the UV–ozone oven was prewarmed for 5 min, the photomask was UV–ozone-cleaned for 10 min and further cleaned with acetone followed by isopropanol, then dried and dusted to ensure good contact. PLL-PEG-coated coverslips were placed (coated side up) on the UV–ozone tray, held in place using vacuum, and the photomask was placed directly on top (chrome side facing the coverslip). Coverslips were UV–ozone-treated for 10 min to locally remove passivation in the mask-defined regions. After patterning, coverslips were either used immediately or stored at 4 °C (patterned side up). Patterned regions were functionalized by coating with fibronectin for 2 h, followed by gentle washing with ultrapure water to remove unbound protein and air-drying. For cell culture, patterned coverslips were mounted into a 4-well chamber, rehydrated for 30 min in water if stored, and cells were seeded onto the patterned areas, allowed to settle briefly, then flooded carefully with culture medium to minimize disturbance. Non-adherent cells were removed by gentle medium exchange once patterns were visible.

##### Micropatterning - Live Cell Experiments

One-well chambered glass slides (Cellvis) were prepared by first attaching a 4-well silicone micro-insert (Ibidi; rectangular well dimensions 2 mm × 1.5 mm) onto the surface of the glass slide. The chambered slide was then plasma-cleaned for 1.5 min and passivated with 10 µL per well of PLL-PEG (0.1 mg mL$^{-1}$, SuSoS) for 1 h at room temperature (22 °C–24 °C). Following passivation, the wells were rinsed 5 times with 10 µL of PBS before adding 10 µL of the PLPP photoinitiator (Alveole) to each well. Disks of varying diameters were then patterned by degradation of the passivation layer with UV light using the PRIMO patterning system (Aveole) at a power setting of 2400 mJ mm$^{-2}$. Immediately following UV patterning, the photoinitiator was removed and the well was rinsed 10 times with 10 µL of PBS. The washed wells were then used for fibronectin coating as described above.

##### Chemical inhibitor treatment

All inhibitors were dissolved in CM at their respective working concentrations and added to dissociated cells after aspiration of DM following centrifugation. Inhibitors remained on the cells for the entire duration of each experiment. When inhibitors were used, the micro-inserts were not filled with CM. Instead, each well was maintained in a separate volume of approximately 15 µL per well, with either inhibitor or vehicle control.

To inhibit actin dynamics and cell movement, a cocktail of three inhibitors, referred to as "JLY", was used: 8 µmol L$^{-1}$ Jasplakinolide (Enzo, ALX-350-275); 1 µmol L$^{-1}$ Latrunculin A (Enzo, BML-T119-0500); and 10 µmol L$^{-1}$ Y-27632 (StemCell Technologies, 72304). Jasplakinolide and Latrunculin A stock solutions were prepared in DMSO, whereas Y-27632 was prepared in phosphate-buffered saline (PBS). Vehicle controls were prepared by adding DMSO to CM to achieve a final concentration of 0.11 % (v/v) DMSO.

To inhibit WNT secretion, 4 µmol L$^{-1}$ IWP-2 (Sigma, I0536) in CM was added to cells after DM was removed. IWP-2 stock solutions were prepared in DMSO, and control conditions contained 0.1 % (v/v) DMSO.

For washout experiments, JLY was removed after 6 h of incubation and wells were washed 5 times with 10 µL of CM. A final 10 µL of CM was added to each well after the last wash, and the entire micro-insert was then filled with 150 µL of CM.

#### Microscopy

Images were acquired on either Zeiss LSM780 or LSM880 laser-scanning confocal microscopes. The image size setting for all experiments ranged from 472 µm to 606 µm. The most common division of the image was $512 \times 512$ pixels with a pixel size of 0.9225 µm. To cover more imaging area in experiments conducted in larger wells, frames were tiled in a non-overlapping manner. To excite and capture eGFP and mCherry fluorescence in cells, a 488 nm laser at 1 % power and a 561 nm laser at 0.5 % power were sequentially scanned using a 1.58 µs pixel dwell time followed by 2- or 4-line averaging. The PMT detector gain was set to 800 or 850 with no digital gain and zero offset. All samples were imaged in an environmental control box heated to 37 °C and supplied with 5 % $CO_2$. For each experiment, 9–25 z-slices with 1 µm to 2 µm spacing were captured at intervals of 2.5 min–5 min per frame for approximately 16 h of total imaging time.

#### Nuclei segmentation and tracking

To segment nuclei, we trained a StarDist 3D model (4) using hand-annotated training data acquired on different microscopes of the same model using similar imaging settings. The StarDist3D model was trained to segment all bright blob-like objects in single-channel H2B-mCherry images. Segmented masks were then processed further to classify objects as debris or cell nuclei. Debris and dying cells were largely characterized by small objects with large nuclear pixel intensity variation. For most analyses, we applied a simple thresholding strategy of 200 µm$^3$ and 0.5 nuclear pixel coefficient of variation to remove debris or dead nuclei. For experiments in large wells in Fig. 3, this simple strategy did not perform as well, likely due to reduced z-sampling. In those cases, we trained a random-forest classifier for each of the three experimental replicates using the Object Classification workflow in Ilastik to label objects as debris or nuclei. Training objects were annotated in Ilastik until acceptable accuracy was achieved by manual inspection.

Segmented nuclei masks from StarDist and raw intensity images were then used as input to the Tracking Workflow in Ilastik. Cell-division and object-count classifiers were trained using between 100 and 1,000 annotated objects for each class. For the object-count classifier, we used three classes: debris, 1 object, and 2 objects. We did not apply the simple thresholding strategy described above for cell tracking, because the object-count classifier already included a debris class. Tracking parameters were left at the suggested default values except for the neighborhood parameter and minimum object size, which were set to 6 and 100, respectively.

#### Experimental *T* spot definition and quantification

For fixed Brachyury immunofluorescence images, each disk was first segmented and the Brachyury channel was binarized using Otsu thresholding. The initial binary mask was cleaned by a morphological opening operation, and connected components smaller than 10% of the area of the smallest disk condition (80 µm disks) were excluded. Spot area was defined as the area of the resulting mask in pixels. Spot diameter was obtained from the segmented spot mask using the minimum Feret diameter returned by the `regionprops` function in MATLAB (2024b).

For live T-reporter imaging, spots were identified manually as local regions of elevated T-reporter signal with an approximately spot-like morphology, based on visual inspection at the 895 minute time point. Center positions were chosen by eye to approximate the center of each visible spot. In the analyses associated with Fig. 1, each apparent spot was counted once and annotated by a manually placed point. For the spot-count analysis used in Fig. 3, manual scoring was performed twice, and the final count was taken using the floor of the two measurements. Nearest-neighbor distances between $T$ spots were computed within each well only, using the manually assigned spot centers.

##### Local correlation between T activity and cell density

To quantify the relationship between local T activity and local cell density, each image was partitioned into a 2D grid of 50 µm × 50 µm bins spanning the image field of view. For each bin, local T activity was defined as the mean per-cell ratio of T-reporter intensity to H2B intensity, where both signals were measured as the mean intensity over nuclear pixels. Local cell density was defined as the number of nuclei whose centroid fell within the same bin. Pearson correlations between local T activity and local cell density were computed separately for each frame and each well using only bins containing more than two cells. The black trace in Fig. 1D represents the mean correlation across wells, while individual well traces are also shown.

A null distribution was obtained by randomly rotating and reflecting the local mean $T$ activity relative to the local cell density 500 times at each time point. Null correlation values were averaged across wells for each randomization to generate the null distribution. The dashed gray line and band in Fig. 1D represent the median and 2.5th–97.5th percentile range of the resulting null distribution.

##### Radial and tangential velocity analysis around aggregation centers

*For Fig. 1E*: Spot centers were defined using the same manually selected $T$ spot annotations. These centers were held fixed in time. Cells were selected for analysis if their tracks lay within 60 µm of a manually selected spot center. The approximately 200 µm × 200 µm inset shown in Fig. 1E is a visualization window for the velocity vectors corresponding to the cells selected for analysis based on the 60 µm criterion.

Radial and tangential velocity components were calculated relative to the fixed spot center using time-averaged cell displacements rather than instantaneous frame-to-frame displacements. Velocities were averaged over the interval from 200 to 590 min post-dissociation. Only tracks with valid positions across the full analysis window were included, because average velocity was defined over the complete interval. Four wells were analyzed in total, with two wells coming from each of two experimental days. Five spots were analyzed per well. On average, more than 50 tracks contributed to each spot-level estimate. Each individual point shown in Fig. 1E represents the average velocity across all included cell tracks for a given spot.

*For Fig. S4E,F*: Cells with valid position measurements across the full interval from 200 to 590 min post-dissociation were assigned at 200 min to the nearest of the five future spot centroids identified for Fig. 1E. Cells were divided into T-high and T-low populations according to whether their T activity (T reporter H2B-EGFP signal divided by H2B-mCherry signal) at 200 minutes was above or below the median within each well. Radial and tangential velocity components were calculated relative to the assigned centroid using the same time-averaged displacement calculation used for the analysis in Fig. 1E, described above. The points shown in Fig. S4E,F represent the resulting well-level means obtained by averaging the five spot-level mean velocities within each well.

A null distribution for the T-high minus T-low velocity difference was obtained by randomly rotating and reflecting the five spot centroids relative to the measured cell tracks 2,000 times within each well. Random rotations differed from the observed orientation by at least 60°. Randomized patterns were retained only if all centroids remained within the image and their mean displacement from the observed centroid positions was at least 60 µm. For each randomization, cells were reassigned to the nearest transformed centroid, and the T-high minus T-low radial and tangential velocity differences were recalculated. Differences were first averaged across spots within each well and then across the four wells as was done in the main analysis. Two-sided p-values were calculated by comparing the absolute deviation of the observed difference from the median of the null distribution with the corresponding absolute deviations of the randomized differences, using a plus-one correction.

##### Fourier-based patterning analysis

Normalized power was used to quantify the strength of patterning over time. It was defined as the sum of Fourier power within the patterning band, $110\,\mu m < \lambda < 250\,\mu m$, divided by the total power in modes with $\lambda < 250\,\mu m$. Fourier power was defined as the squared amplitude of the radially averaged Fourier transform. The wavelength band 110 µm–250 µm was selected because it most closely matched the observed nearest-neighbor spacing distribution of spots and excluded lower-wavenumber structure dominated by the overall well geometry.

This metric was computed primarily from T-reporter images. The main exception was the cross-correlation analysis, where the same metric was also computed from H2B images. Before Fourier analysis, image stacks and the corresponding segmentation masks were loaded. For each channel, the image was smoothed with a Gaussian filter with sigma $[1, 1, 0.5]$, then background-subtracted using the median intensity of non-nuclear pixels. Non-nuclear pixels were defined as pixels outside a nuclear mask dilated by two pixels using a cubic structuring element. Nuclear objects were classified as live cells or debris using a double-threshold rule: live cells were required to have a volume greater than $200\,\mu m^3$ and a coefficient of variation of H2B nuclear intensity less than 0.5. For the JLY experiments, the H2B coefficient-of-variation threshold was relaxed to 0.6 to account for treatment-induced changes in nuclear shape.

After debris exclusion, a zero-valued image of the same size as the original image was created and populated only at voxels belonging to live-cell masks, using the T-reporter signal or, where indicated, the H2B signal. This masked image was then z-scored using only live-cell pixel values, and the resulting image was used for Fourier decomposition.

##### Signal variability and cross-correlation analysis

The coefficient of variation of T expression was computed for each frame from the distribution of T/H2B ratios across all live cells. For the IWP-2 analysis, average T activity was defined as the mean of the T/H2B ratio distribution within each frame.

To compare the timing of patterning and signaling changes, the normalized power and coefficient-of-variation traces were standardized to z-scores separately for each well. For a trace $x_i(t)$ from well $i$, the standardized score was calculated as

$$z_i(t) = \frac{x_i(t) - \overline{x_i}}{s_{x_i}}, \tag{1}$$

where $\overline{x_i}$ and $s_{x_i}$ denote the temporal mean and standard deviation, respectively, of that trace across the analyzed time points. The standardized traces were then subsampled every 10 frames, corresponding to 50 min intervals, differenced, and cross-correlated. The mean cross-correlation across 27 wells is shown in Fig. 2B. Peak lag values were also extracted separately for each well, and the median and interquartile range of those peak lags were used for summary display.

##### Particle Image Velocimetry

The software PIVlab (v. 3.06) within MATLAB (2024b) was used to generate velocity fields using the cell nuclei channel. We used an ensemble three-pass workflow with 128, 64, and 32 pixel window sizes to analyze successive pairs of images in the time-lapse imaging data. The final plotted vector fields represent velocities over frames between 395 min and 595 min post-dissociation. Divergence of the vector field was obtained in a post-processing step within the software. Manually selected aggregation points used for the radial velocity analysis were the same points used in the PIV comparison.

##### Western Blot Analysis

Tail samples were collected from E10.5 embryos by dissection of the PSM, defined as the tissue piece extending from the tip of the tailbud to the first visible somite boundary, in DM. For each genotype, three dissected PSM tissue fragments were rinsed in ice-cold PBS and immediately transferred into 9 µL of ice-cold RIPA buffer for digestion by pipette trituration. Each digested sample was then serially diluted twofold in RIPA buffer, and 2× Laemmli buffer was added to a final volume of 18 µL. All samples were then denatured at 95 °C for 5 min before loading the entire volume onto separate wells of a precast Bio-Rad 4–20% TGX gel. Electrophoresis was run at 20 mA for 1 h with Precision Plus Dual Color Ladder (Bio-Rad, Cat. No. 1610374). After electrophoresis, proteins were transferred from gels to 0.2 µm nitrocellulose membranes (Bio-Rad, Cat. No. 1704158) using the Trans-Blot Turbo system with a 7 min transfer time. Membranes were blocked in 5% milk powder/PBST (0.05% Tween-20) for 1 h and incubated overnight at 4 °C with primary antibodies against Brachyury (R&D, AF2085, 1:1000), GFP (Torrey Pines Biolabs TP401, 1:5000), and HSP90 (Cell Signaling Technology 4874, 1:1000). All primary antibodies were diluted in PBST containing 5% bovine serum albumin and 0.01% sodium azide. Membranes were washed 6–10 times in PBST before secondary antibody incubation with anti-goat-HRP or anti-rabbit HRP at 1:1000 dilution in PBST for 1 h at room temperature. To detect three proteins on the same membrane, the membrane was stripped, blocked, and re-probed after each primary target. Chemiluminescence was detected using SuperSignal West Pico PLUS Substrate (Thermo Fisher 34580) and imaged on a ChemiDoc system (Bio-Rad).

##### Immunofluorescence Staining

*Fixation and staining of samples for size-pattern relationship*: After removal of media, samples were washed with PBS supplemented with calcium and magnesium and then fixed with methanol-DMSO (1:1) for 1 minute at 4 °C. Cells were then washed once with PBS-Triton 1% and blocked for 1 hour at room temperature in a 10% FCS solution with PBS-Triton 1%. Primary antibody against Brachyury (R&D, AF2085) was used at 1:500 dilution and incubated overnight at 4 °C. This was followed by three washes using PBS-Triton 1% followed by incubation with secondary antibody donkey $\alpha$-goat Alexa-Fluor 568 (ThermoFisher A-11057, 1:500) and DAPI (Biotium 40043, 1:1000) overnight at 4 °C.

*Fixation and staining of samples after time-lapse imaging:* Culture medium was aspirated, and samples were washed three times with PBS. Samples were then fixed with cold 4% paraformaldehyde (Electron Microscopy 15710) for 15 min at room temperature. PFA was removed after incubation and samples were washed three times with PBS before a 24 h incubation at 4 °C in blocking and permeabilization buffer (BP) composed of 0.5% Triton X-100 and 10% normal donkey serum in PBS. Primary antibody against Brachyury (R&D, AF2085, 1:20) diluted in BP buffer was then added for another 24 h at 4 °C. Samples were washed three times in PBS before the addition of anti-goat secondary antibody conjugated to Alexa Fluor 647 (Thermo Fisher A-21447, 1:1000) prepared in BP buffer. Secondary antibody incubation was performed at room temperature for 4 h prior to imaging on a Zeiss LSM 880 confocal microscope.

##### Statistical analysis

For analyses of patterning dynamics within circular wells: wells were included only if the average number of cells across all analyzed time points exceeded 800.

For the Fig. 1G cell speed analysis, trajectories were summarized in non-overlapping 5-frame windows, and the reported value was assigned to the center of each window. Because speed did not change appreciably over time, the statistical comparison was based on the average of the first three intervals (15 frames total). Significance was then assessed by comparing treatment differences using a well-level, within-day permutation test.

Linear mixed-effects models throughout the paper were fit in MATLAB (R2024b) using `fitlme` with the formula:

```
Score ~ initial_score + initial_cell_count + drug*time + (1 + time|well)
```

Initial values were computed from the first three frames, and the outcome variable was analyzed from frame 4 onward so that predictor and response windows did not overlap. Significance of the drug $\times$ time interaction effect was then assessed by using well-level, within-day permutation test.

##### Mathematical modeling and analysis

We modeled the distribution of $T+$ cells in space and time $\rho(\mathbf{r}, t)$ using a nonlocal continuum model. The model consists of an advection-diffusion equation, where the local velocity is determined by a weighted integral of cell densities at all positions. On large (infinite) domains, the equation is

$$\partial_t \rho = \nabla \cdot \left( D\nabla\rho - \chi\rho(1 - \rho/\rho_{\max}) \iint_{\mathbb{R}^2} \mathbf{G}(\mathbf{r}' - \mathbf{r})\rho(\mathbf{r}')\mathrm{d}\mathbf{r}' \right). \tag{2}$$

Here $D$ is the effective diffusion constant of cells, accounting for their random motion. The parameter $\chi$ describes the strength of cell-cell interaction and $\mathbf{G}$ is a vector function describing the effect of cells at position $\mathbf{r}'$ on the velocity of cells at position $\mathbf{r}$. The direction of $\mathbf{G}$ is from $\mathbf{r}$ to $\mathbf{r}'$, and its magnitude is a function of the distance $|\mathbf{r}' - \mathbf{r}|$. We use an exponentially decaying interaction strength (see Supplementary theory for the details).

In order to limit the maximal density and model volume exclusion of cells, we adapt the approach of Painter and Hillen (5), also used by Painter (6), who derive volume-filling versions of chemotaxis equations based on a discrete random walk. In their approach, a function $p(\rho)$ is introduced that models how the probability of a cell moving to a new location is modified by the presence of cells at that location. Generically, this function modifies both the advection term and the diffusion term. The advection term is multiplied by $p(\rho)$, whereas the diffusion term is multiplied by $p(\rho) - \rho p'(\rho)$.

We used the simple packing function $p(\rho) = 1 - \rho/\rho_{\max}$. For this linear packing function, $p(\rho) - \rho p'(\rho) = 1$, which means the diffusion term is a constant.

The analysis of this equation leading to the patterning criterion and length scale estimates can be found in Supplemental Theory text.

On bounded domains, we take effects from boundaries into account and modify the equation to

$$\partial_t \rho = \nabla \cdot \Bigg( D\nabla\rho - \rho(1-\rho/\rho_{\max}) \bigg[ \chi \iint_\Omega \mathbf{G}(\mathbf{r}'-\mathbf{r})\rho(\mathbf{r}')\mathrm{d}\mathbf{r}' \\ + \chi_\mathrm{b} \iint_{\mathbb{R}^2 \setminus \Omega} \mathbf{G}_\mathrm{b}(\mathbf{r}'-\mathbf{r})\phi(\mathbf{r}')\mathrm{d}\mathbf{r}' \bigg] \Bigg) . \tag{3}$$

The first interaction kernel, $\mathbf{G}$, describes cell-cell interactions. The integration is only over the region where cells can move: $\Omega$. The second interaction kernel $\mathbf{G}_\mathrm{b}$ describes the effect on cells of the material outside of the system, which is why this integral's domain is everything but $\Omega$, in 2D written as $\mathbb{R}^2\backslash\Omega$. Depending on the system, the boundary material could be external tissues, extracellular matrix, culture medium or something else. The density of boundary material is given by $\phi(\mathbf{r})$, which we assume to be fixed in time. This means that the expression

$$\iint_{\mathbb{R}^2 \setminus \Omega} \mathbf{G}_\mathrm{b}(\mathbf{r}'-\mathbf{r})\phi(\mathbf{r}')\mathrm{d}\mathbf{r}'$$

evaluates to a fixed function of space $\mathbf{B}(\mathbf{r})$, as written in the main text.

The effect of varying $\chi_\mathrm{b}$ was explored in (7). Here, we always use what we have termed *neutral boundary interactions*, which is equivalent to taking a constant outside density $\phi(\mathbf{r}) = \rho_0$, $\mathbf{G}_\mathrm{b} = \mathbf{G}$ and $\chi_\mathrm{b} = \chi$. This choice ensures that the constant density field $\rho(x) = \rho_0$ is a homogeneous steady state of the system.

Eq. (3) is solved on $\Omega$ using no-flux boundary conditions on $\partial\Omega$:

$$D\nabla\rho - \rho(1-\rho/\rho_{\max}) \left[ \chi \iint_\Omega \mathbf{G}(\mathbf{r}'-\mathbf{r})\rho(\mathbf{r}')\mathrm{d}\mathbf{r}' + \chi_\mathrm{b}\rho_0 \iint_{\mathbb{R}^2 \setminus \Omega} \mathbf{G}_\mathrm{b}(\mathbf{r}'-\mathbf{r})\mathrm{d}\mathbf{r}' \right] = 0, \quad \mathbf{r} \in \partial\Omega \tag{4}$$

##### Numerical simulations

To simulate Eq. (3) numerically, we use the method of lines: we first discretize space, leading to a high-dimensional system of ODEs, which we then solve using standard ODE solvers. To discretize space, we use a finite-volume method on a Cartesian grid, which allows for a straightforward implementation of the no-flux boundary. We use a flux limiter (Koren) for the advective flux (8). To compute the integral terms, we use the FFT-based method proposed by Gerisch (9). This method works for periodic domains. To simulate on a bounded domain of arbitrary shape $\Omega$, such as a disk, we embed $\Omega$ in a larger square with periodic boundaries. The velocity field is computed by FFT on the larger square, and density is set to zero outside of $\Omega$. Time-stepping is done using the Runge-Kutta 45 method (`solve_ivp` from `scipy.integrate`). The Python code including demo notebooks can be found at `https://github.com/JanRombouts/zhao-rombouts-scaling`. For long simulations and parameter sweeps we made use of the EMBL High Performance Computing resources (10).

##### Initial conditions

In order to generate initial conditions that approximate the observed density profiles in the beginning of the experiments, we use a method to sample random initial conditions based on the experimental density profiles. We use PCA on the density profiles of $T+$ cells to write the density as a linear combination of orthogonal basis density profiles $\psi_i(\mathbf{r})$. To generate a new density profile, we randomly sample coefficients and recombine them into a density function, i.e. a new initial condition is given by

$$\sum_i \beta_i \psi_i(\mathbf{r}),$$

where $\psi_i(\mathbf{r})$ are the PCA modes from the experiments and $\beta_i$ are randomly sampled coefficients from a normal distribution with mean and standard deviation equal to the mean and standard deviation of the distribution of experimental coefficients. For Fig. 3C, we use the initial density of $T$+ cells from 27 experimental wells with coated diameter of 400 µm. After sampling we rescale the density profile such that the average density is a predetermined number. For Fig. 4 we use the same method based on the 27 wells, followed by a cropping step: we randomly crop out a disk of given size from the large well. A demo notebook detailing the procedure can be found at `https://github.com/JanRombouts/zhao-rombouts-scaling`.

##### Determination of density field from imaging data

To process the imaging data into a density profile of $T$+ cells, we use the following procedure (Fig. S8). After segmentation and removal of dead cells based on the volume and H2B variation in the segmentation mask, we classify each cell as $T$+ or $T$-. We calculate each cell's ratio of T intensity over H2B intensity and assign the 50% highest cells to $T$+. For the micropatterned disks (Fig. 4), smoothing and background subtraction was done on the segmented objects H2B and $T$ channels before removal of dead cells and classification as $T$+/$T$-.

To cancel out the effect of changes in average $T$ intensity, this classification is per frame, i.e. for each frame we take a 50/50 split. From the positions of $T$+ cells we generate a coarse-grained density field. For the large well experiments (Fig. 3B), we simply count the number of cells in each 20 by 20 µm square. For smaller systems, we found that this approach led to obscuring some smaller spatial features. We therefore consider a grid with spacing 5 µm. We defined the density at each grid point by counting the number of cells in a circle of area 400 µm$^2$ centered at the grid point. Before analysis, we temporally smoothed the density profiles with a 11-point running average filter, except for the analysis of the large wells leading to the dispersion relation in Fig. 3.

##### Fitting of parameters of the dispersion relation

We computed the 2D Fourier transform of the density of $T$+ cells at each time point (Fig. S9C). Next, we computed the radial power spectrum by averaging (Fig. S9D). This yields the radial power spectrum $U(k,t)$ where $k$ is the wavenumber. By inspecting the evolution of $U(k,t)$ over time, we identified a time window where the evolution of $U$ is exponential (Fig. S9E). For each of the three experiments we used a 500 min window, but the starting point differed. Performing a linear fit of $\ln U(k,t)$ versus $t$ yields the growth rate $\omega(k)$ (Fig. S9F-H).

To obtain the parameters $D, \chi$ and $\sigma$ from the experimentally determined dispersion relation, we first fit the function (see Supplemental Theory Text for the derivation)

$$f(k) = -Dk^2 + 2\pi\xi\sigma\frac{k^2}{(1+\sigma^2k^2)^{3/2}}$$

to obtain $D$, $\sigma$ and $\xi = \chi\rho_0(1-\rho_0/\rho_{\max})$. The fit was performed using `curve_fit` from the `scipy` library and limited to low wavenumbers, $k < 0.07/(2\pi)$ µm$^{-1}$. This cutoff corresponds to the peaked part of the dispersion relation, excluding the larger wavenumbers (short wavelengths) whose amplitudes seem to be dominated by noise. We determine $\rho_0$ by directly computing the average density over the 500-min interval used. We use $\rho_{\max} = 0.025$ cells/µm$^2$, obtained by inspecting histograms of densities over the different experiments. The code is available at `https://github.com/JanRombouts/zhao-rombouts-scaling`.

For the Bayesian fitting procedure (Fig. S10), we used Approximate Bayesian Computation (11) implemented in the software pyABC (12). We used uniform priors for $D, \sigma, \chi$ and a sum of squares distance function. Hyperparameters of the ABC algorithm can be found in the demo notebook at `https://github.com/JanRombouts/zhao-rombouts-scaling`.

##### Calculation of the dispersion relation on disks

Given neutral boundary conditions, a linear stability analysis (see Supplemental Theory text) on a disk with radius $R$ results in the eigenvalue problem for the growth rate $\omega$, and an eigenmode $u(\mathbf{r})$, with $\mathbf{r} = (r, \theta)$ in polar coordinates,

$$\nabla \cdot (\nabla u - \bar{\alpha}\mathcal{A})\, u = \omega u. \tag{5}$$

The boundary conditions stem from the no-flux requirement

$$[\nabla u - \bar{\alpha}\mathcal{A}]\, u \cdot \hat{\mathbf{n}} = 0, \quad r = R, \tag{6}$$

in which $\hat{\mathbf{n}}$ is the outward pointing normal at the disk boundary. The operator $\mathcal{A}$ is the integral operator

$$\mathcal{A} : f \mapsto \int_\Omega \mathbf{G}(\mathbf{r}' - \mathbf{r}) f(\mathbf{r}') \mathrm{d}\mathbf{r}'. \tag{7}$$

This operator maps a scalar function $f$ on the disk $\Omega$ to a vector function on the disk. We solve the eigenvalue problem numerically in two different ways (the code is available at `https://github.com/JanRombouts/zhao-rombouts-scaling`). First we used a spectral method to discretize the equation in polar coordinates, similar to (13, Chapter 11). In essence, we transform the problem into a generalized eigenvalue problem for matrices:

$$Au = \omega B u,$$

where $B$ is used to encode the boundary conditions.

Since spectral methods can give spurious (non-physical) eigenvalues (14, Sec. 7.6), we also use a more naive discretization based on a Cartesian grid: we divide the disk into pixels and use finite volumes to compute fluxes across each pixel boundary and to implement the boundary conditions. We keep the eigenvalues and modes that are the same in both methods.

##### Simulation spot count determination

To determine the number of spots in the $T+$ density field in simulations, we first used a uniform filter with size 3 on the density field. Next, we used the function `peak_local_max` from `scipy` to determine the peaks of the density profile. We used a minimum distance between peaks of 15 grid points and minimum peak value of $\rho_{\max}/2$. In addition we excluded peaks closer than $R/20$ to the boundary of disks, where $R$ is the disk radius. The GitHub repository contains a notebook with a demonstration of how we analyze simulation output.

##### Analysis of temporal evolution of patterns on small disks

We obtain the density of $T+$ cells in a similar way as for the larger circular wells. To avoid an excessive influence of sparse cells near the borders of the disks, we keep only grid points for which the circle in which cells are counted lies at least for 75% inside the disk domain. We consider the time interval between 200 and 1000 minutes post tissue dissociation.

We discard disks for which the average density over this interval is less than 0.005 cells/$\mu m^2$, or for which the density change (calculated as the maximal minus the minimal value) is larger than 0.005 cells/$\mu m^2$.

We calculate the normalized eigenmodes from the linear stability analysis for each disk size separately and expand the density profile for each timepoint into the first 7 modes (excluding the radially symmetric mode with growth rate 0), after subtracting the mean density. This set of modes includes polar, central-spot, bipolar and tripolar modes. We thus have

$$\rho(\mathbf{r}, t) = \rho_0(t) + \sum_{i=0}^{6} a_i(t)\phi_i(\mathbf{r}).$$

Each non-radially symmetric mode appears twice, with two different orientations. We combine coefficients for modes that are rotated versions of each other using root-mean-squares to obtain mode amplitudes. The dominant mode at a timepoint is defined as the one with maximal amplitude at that timepoint. The expansion procedure can be found in a demo notebook at `https://github.com/JanRombouts/zhao-rombouts-scaling`.

##### Determination of scaling regime

To determine the scaling regime in Fig. 4F, we performed 100 simulations for each system size. For each simulation, we detected the time at which the system shows a single-spot pattern based on a decomposition in Bessel modes. We classify the pattern as single spot when the total power in the modes with angular/radial numbers $(0, 1)$ (center spot) and $(1, 1)$ (polarized spot) divided by the total power in all modes is larger than 0.4. The scaling regime is defined as the region where at least half of the simulations exhibit single spot outcome. The code is available at `https://github.com/JanRombouts/zhao-rombouts-scaling`.

### Supplemental theory

Throughout we use bold notation for vectorial quantities.

#### Model equations and scaling

We use the advection-diffusion equation

$$\partial_t \rho = \nabla \cdot (D\nabla\rho - \mathbf{v}\rho) \tag{8}$$

to describe the time evolution of the density of $T+$ cells on a space $\Omega$. The number of cells is conserved. The velocity at location $\mathbf{r}$ is determined by the density distribution of cells, and by influences from material outside the domain:

$$\mathbf{v}(\mathbf{r}) = p(\rho)\left[\chi \iint_\Omega \mathbf{G}(\mathbf{r}'-\mathbf{r})\rho(\mathbf{r}')\mathrm{d}\mathbf{r}' + \chi_\mathrm{b}\mathbf{B}(\mathbf{r})\right] \tag{9}$$

In this expression, $p(\rho)$ is called a *packing function*, which limits the velocity for high densities. It can be derived from random-walk models where the jumping probability depends on the occupation of a target site (5). In this work, we use

$$p(\rho) = 1 - \rho/\rho_\mathrm{max}. \tag{10}$$

The integral expression describes cell-cell interactions, with the function $\mathbf{G}$ acting as the interaction kernel that describes the direction and strength of the interaction as a function of relative cell position $\mathbf{r}'-\mathbf{r}$. We use the expression

$$\mathbf{G}(\mathbf{r}'-\mathbf{r}) = \frac{1}{\sigma^2} g(|\mathbf{r}'-\mathbf{r}|/\sigma)\frac{\mathbf{r}'-\mathbf{r}}{|\mathbf{r}'-\mathbf{r}|}. \tag{11}$$

Here $\sigma$ is the characteristic length scale of the interaction, and we use an exponentially decaying interaction strength $g(r) = e^{-r}$. The vectorial part $(\mathbf{r}'-\mathbf{r})/|\mathbf{r}'-\mathbf{r}|$ is the unit vector pointing from $\mathbf{r}$ to $\mathbf{r}'$. This functional form can be derived from mechanistic PDE models in simplified settings (7, 15).

In principle, the boundary effect $\mathbf{B}(\mathbf{r})$ can be any function, but we also model it using an interaction kernel:

$$\mathbf{B}(\mathbf{r}) = \int_{\mathbb{R}^2\setminus\Omega} \mathbf{G}_\mathrm{b}(\mathbf{r}'-\mathbf{r})\phi(\mathbf{r}')\mathrm{d}\mathbf{r}'. \tag{12}$$

The density field $\phi$ describes the density of outside material. For simplicity, in this work we use $\mathbf{G}_\mathrm{b} = \mathbf{G}$. Additionally, we write $\phi(\mathbf{r}) = \phi_0$, such that the density of outside material is constant.

In full, the equation thus reads

$$\partial_t\rho = \nabla\cdot\left(D\nabla\rho - \rho(1-\rho/\rho_\mathrm{max})\left[\chi\iint_\Omega \mathbf{G}(\mathbf{r}'-\mathbf{r})\rho(\mathbf{r}')\mathrm{d}\mathbf{r}' + \chi_\mathrm{b}\phi_0\iint_{\mathbb{R}\setminus\Omega}\mathbf{G}(\mathbf{r}'-\mathbf{r})\mathrm{d}\mathbf{r}'\right]\right). \tag{13}$$

We scale time, space and the density as follows:

$$\hat{t} = tD/\sigma^2, \quad \hat{\mathbf{r}} = \mathbf{r}/\sigma, \quad \hat{\rho}(\hat{t},\hat{\mathbf{r}}) = \rho(t,\mathbf{r})/\rho_\mathrm{max}.$$

Now,

$$\nabla_\mathbf{r} = \nabla_{\hat{\mathbf{r}}}\frac{1}{\sigma}, \quad \partial_t = \partial_{\hat{t}} D/\sigma^2$$

and the nonlocal term becomes

$$
\begin{aligned}
\iint_{\Omega} \mathbf{G}(\mathbf{r}'-\mathbf{r})\rho(\mathbf{r}') &= \iint_{\Omega} \frac{1}{\sigma^2} g(|\mathbf{r}'-\mathbf{r}|/\sigma) \frac{\mathbf{r}'-\mathbf{r}}{|\mathbf{r}'-\mathbf{r}|}\rho(\mathbf{r}')\mathrm{d}\mathbf{r}' \\
&= \iint_{\hat{\Omega}} \frac{1}{\sigma^2} g(|\hat{\mathbf{r}'}-\hat{\mathbf{r}}|) \frac{\hat{\mathbf{r}'}-\hat{\mathbf{r}}}{|\hat{\mathbf{r}'}-\hat{\mathbf{r}}|}\rho(\sigma\hat{\mathbf{r}'})\sigma^2\mathrm{d}\hat{\mathbf{r}'} \qquad \hat{\mathbf{r}'} = \mathbf{r}'/\sigma, \hat{\mathbf{r}} = \mathbf{r}/\sigma \\
&= \iint_{\hat{\Omega}} g(|\hat{\mathbf{r}'}-\hat{\mathbf{r}}|) \frac{\hat{\mathbf{r}'}-\hat{\mathbf{r}}}{|\hat{\mathbf{r}'}-\hat{\mathbf{r}}|}\rho_{\max}\hat{\rho}(\hat{\mathbf{r}'})\mathrm{d}\hat{\mathbf{r}'}
\end{aligned}
$$

The integral in the boundary term, similarly, becomes

$$
\iint_{\mathbb{R}^2\backslash\Omega} \mathbf{G}(\mathbf{r}'-\mathbf{r})\mathrm{d}\mathbf{r}' = \iint_{\mathbb{R}^2\backslash\hat{\Omega}} g(|\hat{\mathbf{r}'}-\hat{\mathbf{r}}|)\frac{\hat{\mathbf{r}'}-\hat{\mathbf{r}}}{|\hat{\mathbf{r}'}-\hat{\mathbf{r}}|}\mathrm{d}\hat{\mathbf{r}'}.
$$

Note that the domain changes from $\Omega$ to $\hat{\Omega}$.

Plugging everything in in Eq. (13) and dropping the hats for clarity gives

$$
\partial_t \rho = \nabla \cdot \left( \nabla\rho - \rho(1-\rho) \left[ \alpha \iint_{\Omega} g(|\mathbf{r}'-\mathbf{r}|)\frac{\mathbf{r}'-\mathbf{r}}{|\mathbf{r}'-\mathbf{r}|}\rho(\mathbf{r}')\mathrm{d}\mathbf{r}' + \beta \iint_{\mathbb{R}^2\backslash\Omega} g(|\mathbf{r}'-\mathbf{r}|)\frac{\mathbf{r}'-\mathbf{r}}{|\mathbf{r}'-\mathbf{r}|}\mathrm{d}\mathbf{r}' \right] \right) \tag{14}
$$

where

$$
\alpha = \frac{\chi\sigma\rho_{\max}}{D} \qquad \text{and} \qquad \beta = \frac{\chi_{\mathrm{b}}\sigma\phi_0}{D}.
$$

The dynamics of this equation depend on parameter $\alpha$, which describes the strength of cell-cell interaction compared to random motion, on $\beta$, which describes cell-boundary interactions, and on the system on which $\rho$ evolves, $\Omega$. First, we will analyze an infinite system where $\Omega = \mathbb{R}^2$. In this case, there is no boundary term.

#### Linear stability analysis on an infinite domain

We analyze the equation with $\Omega = \mathbb{R}^2$ and no boundary term:

$$
\partial_t \rho = \nabla \cdot \left( \nabla\rho - \alpha\rho(1-\rho) \iint_{\mathbb{R}^2} g(|\mathbf{r}'-\mathbf{r}|)\frac{\mathbf{r}'-\mathbf{r}}{|\mathbf{r}'-\mathbf{r}|}\rho(\mathbf{r}')\mathrm{d}\mathbf{r}' \right). \tag{15}
$$

On an infinite domain, we have translation invariance. This entails that

$$
\begin{aligned}
\iint_{\mathbb{R}^2} g(|\mathbf{r}'-\mathbf{r}|)\frac{\mathbf{r}'-\mathbf{r}}{|\mathbf{r}'-\mathbf{r}|}\mathrm{d}\mathbf{r}' &= \iint_{\mathbb{R}^2} g(|\mathbf{s}|)\frac{\mathbf{s}}{|\mathbf{s}|}\mathrm{d}\mathbf{s} \\
&= \int_0^{2\pi}\int_0^{\infty} g(r)\begin{pmatrix}\cos\theta \\ \sin\theta\end{pmatrix} r\mathrm{d}r\mathrm{d}\theta \\
&= \mathbf{0}.
\end{aligned}
$$

Essentially, this states that in a homogeneous density field, all interaction contributions at a certain location will cancel out, leading to a net zero velocity. This entails that any constant density profile $\rho(\mathbf{r}) = \rho_0$ is a spatially homogeneous steady state of Eq. (15).

To assess whether self-organized patterns can appear around $\rho(\mathbf{r}) = \rho_0$, we write $\rho(\mathbf{r}, t) = \rho_0 + \tilde{\rho}(\mathbf{r}, t)$, plug this into Eq. (15) and keep only terms linear in $\tilde{\rho}$. This gives

$$\partial_t \tilde{\rho} = \nabla^2 \tilde{\rho} - \bar{\alpha} \nabla \cdot \iint_{\mathbb{R}^2} g(|\mathbf{r}' - \mathbf{r}|) \frac{\mathbf{r}' - \mathbf{r}}{|\mathbf{r}' - \mathbf{r}|} \tilde{\rho}(\mathbf{r}') \mathrm{d}\mathbf{r}', \tag{16}$$

where

$$\bar{\alpha} = \alpha \rho_0 (1 - \rho_0). \tag{17}$$

On infinite domains, solutions to this linear equation are combinations of functions of the form $e^{\omega t + i\mathbf{k}\cdot\mathbf{r}}$, that describe the time evolution of a spatial mode with wavenumbers $\mathbf{k} = (k_x, k_y)$ and using $\mathbf{r} = (x, y)$. Substituting this form into Eq. (16) will yield a relation between $\omega$ and $|\mathbf{k}|$. Before we do this, we perform some preliminary calculations. We know that

$$\nabla^2 e^{\omega t + i k_x x + i k_y y} = -(k_x^2 + k_y^2) e^{\omega t + i k_x x + i k_y y} = -|\mathbf{k}|^2 e^{\omega t + i\mathbf{k}\cdot\mathbf{r}},$$

but a bit of work is needed to compute the effect of the nonlocal term on this function. We have

$$\begin{aligned}
\nabla \cdot \left( \iint_{\mathbb{R}^2} g(|\mathbf{r}' - \mathbf{r}|) \frac{\mathbf{r}' - \mathbf{r}}{|\mathbf{r}' - \mathbf{r}|} e^{i\mathbf{k}\cdot\mathbf{r}'} d\mathbf{r}' \right) &= \nabla \cdot \left( \iint_{\mathbb{R}^2} g(|\mathbf{s}|) \frac{\mathbf{s}}{|\mathbf{s}|} e^{i\mathbf{k}\cdot(\mathbf{r}+\mathbf{s})} d\mathbf{s} \right) \\
&= \nabla \cdot \left( \underbrace{e^{i\mathbf{k}\cdot\mathbf{r}}}_{\text{scalar function of } \mathbf{r}} \underbrace{\iint_{\mathbb{R}^2} g(|\mathbf{s}|) \frac{\mathbf{s}}{|\mathbf{s}|} e^{i\mathbf{k}\cdot\mathbf{s}} \mathrm{d}\mathbf{s}}_{\text{constant vector}} \right).
\end{aligned} \tag{18}$$

For scalar functions $f(\mathbf{r})$ and a constant vector $\mathbf{v}$, we know (using for example the product rule for the $\nabla$ operator) that $\nabla \cdot (f(\mathbf{r})\mathbf{v}) = (\nabla f) \cdot \mathbf{v}$. Since $\nabla(e^{i\mathbf{k}\cdot\mathbf{r}}) = i\mathbf{k} e^{i\mathbf{k}\cdot\mathbf{r}}$, Eq. (18) equals

$$i e^{i\mathbf{k}\cdot\mathbf{r}} \left( \mathbf{k} \cdot \iint_{\mathbb{R}^2} g(|\mathbf{s}|) \frac{\mathbf{s}}{|\mathbf{s}|} e^{i\mathbf{k}\cdot\mathbf{s}} \mathrm{d}\mathbf{s} \right). \tag{19}$$

We now plug in $\tilde{\rho}(\mathbf{r}, t) = e^{\omega t + i\mathbf{k}\cdot\mathbf{r}}$ into Eq. (16), use the calculations above to work out the algebra, to find

$$\omega = -|\mathbf{k}|^2 - \bar{\alpha} i \left( \mathbf{k} \cdot \iint_{\mathbb{R}^2} g(|\mathbf{s}|) \frac{\mathbf{s}}{|\mathbf{s}|} e^{i\mathbf{k}\cdot\mathbf{s}} \mathrm{d}\mathbf{s} \right). \tag{20}$$

The two-dimensional transform can be simplified further (16). We have that

$$\iint_{\mathbb{R}^2} g(|\mathbf{s}|) \frac{\mathbf{s}}{|\mathbf{s}|} e^{i\mathbf{k}\cdot\mathbf{s}} \mathrm{d}\mathbf{s} = -i \nabla_{\mathbf{k}} \left( \iint_{\mathbb{R}^2} \frac{g(|\mathbf{s}|)}{|\mathbf{s}|} e^{i\mathbf{k}\cdot\mathbf{s}} \mathrm{d}\mathbf{s} \right) = -i \nabla_{\mathbf{k}} \left( 2\pi \int_0^\infty g(s) J_0(ks) \mathrm{d}s \right),$$

where we write $s = |\mathbf{s}|$ and $k = |\mathbf{k}|$. The last equality comes from (16, Eq. 7). It is the 2D Fourier transform of the isotropic function $y \mapsto g(y)/y$. Here $J_0$ is the Bessel function of the first kind of order 0. Further, we have

$$\nabla_{\mathbf{k}} J_0(ks) = \left( \partial_{k_x} J_0 \left( \sqrt{k_x^2 + k_y^2} s \right), \partial_{k_y} J_0 \left( \sqrt{k_x^2 + k_y^2} s \right) \right) = \frac{\mathbf{k}}{k} \frac{d}{dk} J_0(ks)$$

such that

$$i \left( \mathbf{k} \cdot \iint_{\mathbb{R}^2} g(|\mathbf{s}|) \frac{\mathbf{s}}{|\mathbf{s}|} e^{i\mathbf{k}\cdot\mathbf{s}} \mathrm{d}\mathbf{s} \right) = i\mathbf{k} \cdot (-i) \frac{\mathbf{k}}{k} 2\pi \int_0^\infty \mathrm{d}s g(s) \frac{d}{dk} J_0(ks) = 2\pi k \int_0^\infty s g(s) J_0'(ks) \mathrm{d}s.$$

Now, using $J_0'(ks) = -J_1(ks)$, we can write the dispersion relation:

$$\omega(k) = -k^2 + 2\pi \bar{\alpha} k \int_0^\infty g(s) s J_1(ks) \mathrm{d}s = -k^2 + 2\pi \bar{\alpha} k \hat{g}^H(k), \tag{21}$$

where

$$\hat{g}^H(k) = \int_0^\infty g(s)sJ_1(ks)\mathrm{d}s$$

is the Hankel transform of the function $g$. This dispersion relation is still valid for any interaction function $g$. In our study, we use $g(r) = e^{-r}$ , for which the Hankel transform is

$$\frac{k}{(1+k^2)^{3/2}}.$$

We thus, finally, find

$$\omega(k) = -k^2 + 2\pi\bar{\alpha}\frac{k^2}{(1+k^2)^{3/2}}. \tag{22}$$

The state $\rho(\mathbf{r}) = \rho_0$ is linearly stable if $\omega(k) < 0$ for all $k$, and unstable when there exist $k$ such that $\omega(k) > 0$. Whether this is true depends on the value of $\bar{\alpha}$ (Fig. S9A). There is a threshold value of $\bar{\alpha}$, above which the curve has a maximum for $k > 0$. There are different ways to show this threshold is $1/(2\pi)$. The following is a direct algebraic/graphical solution. First, write $q = k^2$ such that the dispersion relation is

$$\omega(q) = -q + 2\pi\bar{\alpha}q/(1+q)^{3/2}.$$

A maximum requires $\frac{d\omega}{dq} = 0$, which means

$$1 - 2\pi\bar{\alpha}\frac{1-q/2}{(1+q)^{5/2}} = 0,$$

or equivalently

$$2\pi\bar{\alpha}(1-q/2) = (1+q)^{5/2}. \tag{23}$$

Sketching both sides of this equation as a function of $q$ shows that an intersection is only possible if the intersection of the graph of $2\pi\bar{\alpha}(1-q/2)$ with the vertical axis has a $y$-coordinate larger than one. This, in turn, requires $2\pi\bar{\alpha} = 1$ or

$$\bar{\alpha} = \frac{1}{2\pi}, \tag{24}$$

which is indeed the instability threshold.

#### Dispersion relation with physical units

Eq. (22) has the nondimensional growth rate and wavelength. We transform back to dimensional units using the timescale $\sigma^2/D$ and space scale $\sigma$. This yields ($\omega$ and $k$ now *with* units despite being denoted by the same symbol as before).

$$\begin{aligned}\omega &= -Dk^2 + 2\pi\bar{\alpha}D\frac{k^2}{(1+\sigma^2k^2)^{3/2}} \\ &= -Dk^2 + 2\pi\chi\rho_{\max}\sigma\rho_0(1-\rho_0)\frac{k^2}{(1+\sigma^2k^2)^{3/2}}.\end{aligned} \tag{25}$$

In this last equation, $\rho_0$ is still the scaled density, i.e. measured in units of $\rho_{\max}$.

#### Estimate of the wavelength and growth rate from the dispersion relation

The wavenumber $k$ with the largest growth rate will dominate in the early stages of patterning. Since a wavenumber $k$ corresponds to the spatial mode $e^{i\mathbf{k}\cdot\mathbf{r}}$ with $|\mathbf{k}| = k$, by knowing this dominant wavenumber we can extract a value of the dominating wavelength of the (repeating) pattern. The dominant wavenumber solves $\frac{d\omega}{dk} = 0$, which as we have seen above leads to

$$2\pi\bar{\alpha}(1 - q/2) = (1 + q)^{5/2}, \tag{26}$$

with $q = k^2$. Generally, we cannot find an analytical solution to this equation, but we can obtain an approximation. Since we are mostly interested in the patterning regime, we look at an approximation for $\bar{\alpha}$ large. Set $\epsilon = 1/\bar{\alpha}$. We will now analyze the solution of

$$2\pi(1 - q/2) = \epsilon(1 + q)^{5/2} \tag{27}$$

for $\epsilon \to 0$. We take the ansatz $q = q_0 + \epsilon q_1$, plug this into the equation and solve the resulting equations order by order. The order 1 equation is

$$2\pi(1 - q_0/2) = 0,$$

which immediately yields $q_0 = 2$. The equation of order $\epsilon$ is

$$-\pi q_1 = 3^{5/2} \tag{28}$$

from which $q_1 = -3^{5/2}/\pi$ and we find the approximation

$$q = 2 - \frac{3^{5/2}}{\pi\bar{\alpha}} + \mathcal{O}(1/\bar{\alpha}^2) \tag{29}$$

The wavelength itself is

$$\lambda = \frac{2\pi}{k} = \frac{2\pi}{q^{1/2}} \approx 2\pi\left(2 - \frac{3^{5/2}}{\pi\bar{\alpha}}\right)^{-1/2},$$

which we can also expand up to first order in $1/\bar{\alpha}$. This gives

$$\lambda \approx 2\pi 2^{-1/2}\left(1 + \frac{1}{4}\frac{3^{5/2}}{\pi\bar{\alpha}}\right) = \pi 2^{1/2} + \frac{2^{1/2}3^{5/2}}{4}\frac{1}{\bar{\alpha}} \tag{30}$$

This wavelength is in dimensionless units, we multiply by $\sigma$ to obtain the unscaled wavelength, and also substitute $\bar{\alpha}$, to find

$$\lambda = A\sigma + B\frac{D}{\chi\rho_{\max}}\frac{1}{\rho_0(1-\rho_0)}, \tag{31}$$

where $\rho_0$ is still the scaled density (i.e. the density relative to $\rho_{\max}$) and

$$A = \pi 2^{1/2} \approx 4.44 \quad \text{and} \quad B = \frac{2^{1/2}3^{5/2}}{4} \approx 5.51. \tag{32}$$

This shows that the wavelength of the pattern is dominated by a term proportional to $\sigma$, with a correction due to $D, \chi$ and the density. In particular, for scaled densities smaller than 1/2, which is the regime our experiments fall into, equation Eq. (31) shows that the wavelength decreases with increasing $\rho_0$. Since the wavelength is roughly the distance between spots, this entails increasing $\rho_0$ is expected to correspond to more spots on a given system size.

To turn this dependence of wavelength into an estimate of the spot count, as shown in Fig. 3C in the main text, we use the following approximation. Since the wavelength $\lambda$ roughly corresponds to the distance between two spots, we assume that each spot including empty space around it takes up the area of a circle with radius $\lambda/2$. With $\lambda$ given by Eq. (31), we then calculate the number of spots the well could accommodate by dividing the total well area by the area of a circle with radius $\lambda/2$.

The growth rate associated with this maximal wavelength can be obtained by plugging the values of $q$ found in Eq. (29) into the original dispersion relation and expanding for small $\epsilon$ (large $\bar{\alpha}$). Eventually, we obtain

$$\begin{aligned}\omega &= \frac{2\pi q_0}{(1+q_0)^{3/2}}\bar{\alpha} + 2\pi\frac{q_1}{(1+q_0)^{3/2}}\left(1-\frac{3}{2}\frac{q_0}{1+q_0}\right) - q_0 + \mathcal{O}(1/\bar{\alpha}) \\ &= 4\pi 3^{-3/2}\bar{\alpha} - 2 + \mathcal{O}(1/\bar{\alpha}).\end{aligned}$$

To obtain the growth rate in physical units, we multiply by the inverse timescale $D/\sigma^2$, and finally find that

$$\omega \approx K\frac{\chi\rho_{\max}}{\sigma}\rho_0(1-\rho_0) + L\frac{D}{\sigma^2}, \qquad (33)$$

with

$$K = 4\pi 3^{-3/2} \approx 2.42 \quad \text{and} \quad L = -2$$

and $\rho_0$ in units of $\rho_{\max}$. In particular, the leading term in the approximation scales inversely with $\sigma$.

These formulae hold for large systems, where boundaries do not play an important role. In addition, since they are derived from the linear stability analysis, these estimates are valid in the initial stages of patterning. Later we consider the linear stability analysis for smaller systems.

#### Estimate of a characteristic velocity

In the main text we claim that $\chi\rho$ is a typical value of the velocity. Here, we substantiate this claim and show that, in the model, the maximal velocity is $2\chi\rho$.

In the scaled equation Eq. (14), the velocity field (assuming no boundary interactions) is given by

$$\mathbf{v}(\mathbf{r}) = (1-\rho)\alpha\iint_{\mathbb{R}^2} g(|\mathbf{r}'-\mathbf{r}|)\frac{\mathbf{r}'-\mathbf{r}}{|\mathbf{r}'-\mathbf{r}|}\rho(\mathbf{r}')\mathrm{d}\mathbf{r}'. \qquad (34)$$

We consider the situation of a density field $\rho$ that is a constant $\bar{\rho}$ in a region $A$ and zero elsewhere, and look at the velocity at a point $\mathbf{r}$ with zero packing, i.e. cells there can move unencumbered. In that case,

$$\mathbf{v}(\mathbf{r}) = \alpha\bar{\rho}\iint_A g(|\mathbf{r}'-\mathbf{r}|)\frac{\mathbf{r}'-\mathbf{r}}{|\mathbf{r}'-\mathbf{r}|}\mathrm{d}\mathbf{r}', \qquad (35)$$

and we only need to investigate which values the integral can take. This is generally hard to do analytically, so we consider a special case and calculate the maximal value of this integral. This is obtained when $A$ is a half plane with $\mathbf{r}$ on the edge (Fig. S11A). After choosing a coordinate system, the velocity will only have an $x$-component. For our typical kernel $g(r) = e^{-r}$, the velocity in the $x$-direction is given by

$$\begin{aligned}\int_{-\pi/2}^{\pi/2}\mathrm{d}\theta\int_0^\infty r\cos\theta e^{-r}\mathrm{d}r &= \int_{-\pi/2}^{\pi/2}\mathrm{d}\theta\cos\theta\int_0^\infty re^{-r}\mathrm{d}r \\ &= 2.\end{aligned}$$

We thus have for the scaled speed $v = 2\alpha\bar{\rho}$. Plugging in $\alpha = \chi\sigma\rho_{\max}/D$ and multiplying by the length scale $\sigma$ and dividing by the time scale $\sigma^2/D$ gives the estimate $v = 2\chi\rho_{\max}\bar{\rho}$, where, since $\bar{\rho}$ was in scaled units, this is thus $2\chi$ times a typical density. Arguably, the situation here gives the maximal value the velocity field can take due to a constant density.

As a second example, we consider the situation shown in Fig. S11B-C: a disk of radius $R$ with constant density $\bar{\rho}$, and we wish to know the velocity field at a distance $r_0$ from this disk. We could interpret this as the velocity of a cell moving towards a spot of $T+$ cells. We numerically compute the required integral and find that it decreases (roughly exponentially) with $r_0$, and is larger as $R$ is larger. The limit $r_0 = 0$ and $R \to \infty$ correspond to the half plane, for which the integral is 2 as shown above.

#### Linear stability analysis on a bounded domain

We start from the equation with a boundary term, in non-dimensional form (Eq. (14)):

$$\partial_t \rho = \nabla \cdot \left( \nabla \rho - \rho(1-\rho) \left[ \alpha \iint_\Omega g(|\mathbf{r}' - \mathbf{r}|) \frac{\mathbf{r}' - \mathbf{r}}{|\mathbf{r}' - \mathbf{r}|} \rho(\mathbf{r}') \mathrm{d}\mathbf{r}' + \beta \iint_{\mathbb{R}^2 \setminus \Omega} g(|\mathbf{r}' - \mathbf{r}|) \frac{\mathbf{r}' - \mathbf{r}}{|\mathbf{r}' - \mathbf{r}|} \mathrm{d}\mathbf{r}' \right] \right) \quad (36)$$

with zero-flux boundary condition

$$\nabla \rho - \rho(1-\rho) \left[ \alpha \iint_\Omega g(|\mathbf{r}' - \mathbf{r}|) \frac{\mathbf{r}' - \mathbf{r}}{|\mathbf{r}' - \mathbf{r}|} \rho(\mathbf{r}') \mathrm{d}\mathbf{r}' + \beta \iint_{\mathbb{R}^2 \setminus \Omega} g(|\mathbf{r}' - \mathbf{r}|) \frac{\mathbf{r}' - \mathbf{r}}{|\mathbf{r}' - \mathbf{r}|} \mathrm{d}\mathbf{r}' \right] = 0, \quad \mathbf{r} \in \partial\Omega \quad (37)$$

Recall that

$$\alpha = \frac{\chi \sigma \rho_{\max}}{D} \quad \text{and} \quad \beta = \frac{\chi_{\mathrm{b}} \sigma \phi_0}{D}$$

are the dimensionless parameters describing cell-cell and cell-boundary material interactions respectively.

We use the *neutral boundary interaction*, $\beta = \alpha \rho_0$ such that the uniform density profile $\rho(\mathbf{r}) = \rho_0$ is a steady state of the system. Setting $\rho = \rho_0 + \tilde{\rho}(\mathbf{r}, t)$, the linearized equation is

$$\partial_t \tilde{\rho} = \nabla^2 \tilde{\rho} - \bar{\alpha} \nabla \cdot \iint_\Omega g(|\mathbf{r}' - \mathbf{r}|) \frac{\mathbf{r}' - \mathbf{r}}{|\mathbf{r}' - \mathbf{r}|} \tilde{\rho}(\mathbf{r}') \mathrm{d}\mathbf{r}', \quad (38)$$

with $\bar{\alpha} = \alpha \rho_0 (1 - \rho_0)$. This equation is identical to Eq. (16), except for the integration domain. In contrast to the infinite-domain case, functions of the form $e^{i\mathbf{k}\cdot\mathbf{r}}$ are no longer the correct spatial modes to use, since they are not eigenfunctions of the spatial operator. Rather, we look for solutions to Eq. (38) of the form

$$\tilde{\rho}(\mathbf{r}, t) = u(\mathbf{r}) e^{\omega t},$$

which after substitution into the equation yields the eigenvalue problem

$$(\nabla^2 - \bar{\alpha} \nabla \cdot \mathcal{A}) u = \omega u. \quad (39)$$

The operator $\mathcal{A}$ is the integral operator

$$\mathcal{A} : f \mapsto \int_\Omega \mathbf{G}(\mathbf{r}' - \mathbf{r}) f(\mathbf{r}') \mathrm{d}\mathbf{r}'. \quad (40)$$

Note that it maps a scalar function $f$ on the disk $\Omega$ to a vector function on the disk. Eq. (39) is complemented by the zero-flux boundary conditions on u

$$(\nabla - \bar{\alpha} \mathcal{A}) u \cdot \hat{\mathbf{n}} = 0 \quad \text{on } \partial\Omega, \quad (41)$$

meaning the radial component of $(\nabla u - \bar{\alpha} \mathcal{A}) u$ should be zero on the boundary of the disk. The solutions $\omega$, $u$ of this equation give the growth rates and eigenmodes. They depend on the parameter $\bar{\alpha}$, as well as on the disk size. We computed the solutions numerically (see Methods).

### Supplemental figures

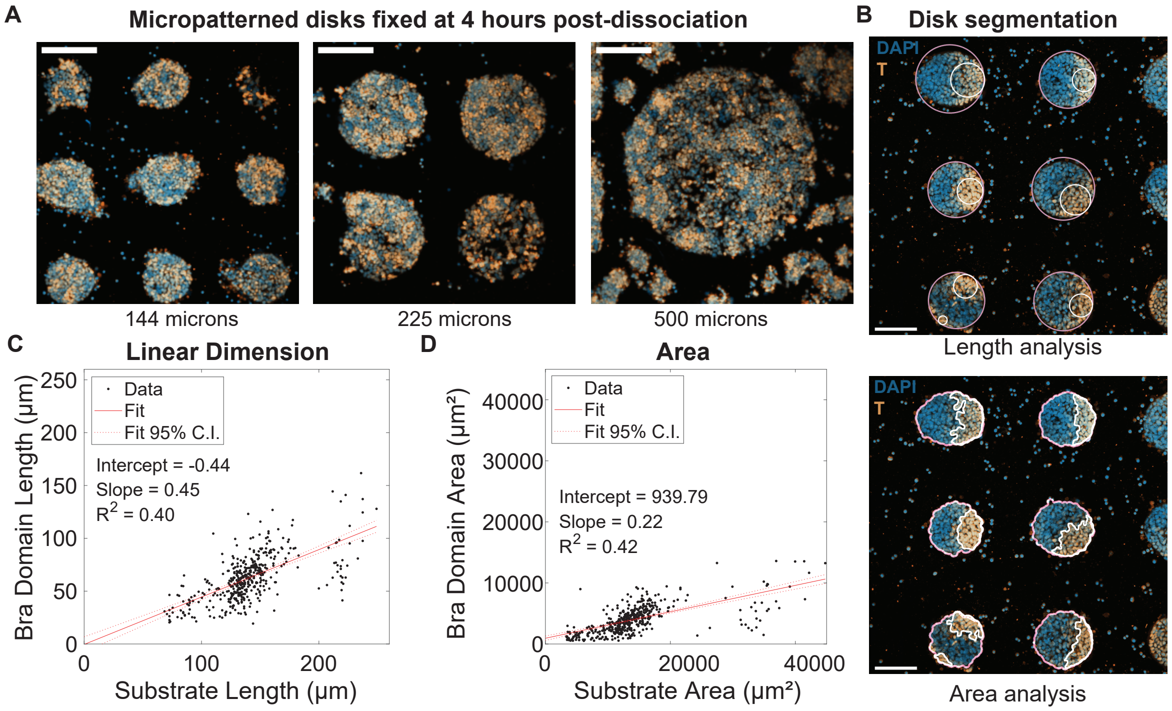


Figure S1: **Analysis of Brachyury patterning on disks of varying diameters A)** Examples of disks fixed 4 hours post-dissociation. Images are composites of nuclei (blue) and Brachyury (yellow) staining. Three disks diameters are shown from left to right: 144, 225, and 500 µm. Scale bar: 150 µm. **B)** Example outputs from automated disk analysis. The top image is an example of a measurement of the linear dimension using bounding circles of the nuclei region and of the Brachyury expression domain. The bottom image represents the segmentation outline of the nuclei region and Brachyury domains used to quantify area. Scale bar: 100 µm. **C)** The correlation between the linear dimension of the disk substrate and the Brachyury domain. **D)** The correlation between the areas of the disk substrate and the Brachyury domain. Only disks with single spot patterns were analyzed, and actual disk sizes were determined by quantifying the area covered by the nuclei staining.

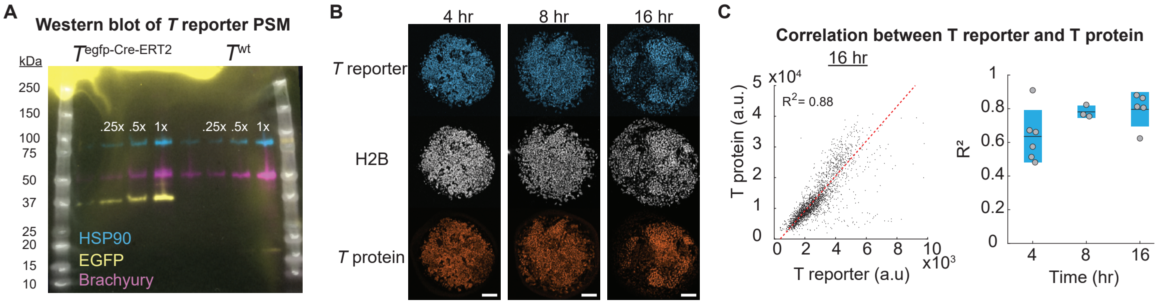


Figure S2: **Validation of Brachyury reporter A)** Western Blots against Brachyury and GFP with HSP90 as the loading control. The composite image was formed from 3 sequential detections using the ladder as the point of reference for alignment. No obvious ectopic fusion protein bands were detected under these conditions. **B)** Spatial comparisons of the $T$ reporter and $T$ protein levels across different time points during ePSM patterning. Time is relative to tissue dissociation as described in Fig. 1A. Scale bar: 100 µm. **C)** Single cell correlation between $T$ reporter and $T$ protein for the three time points in B). The left scatter plot is one example of the correlation at 16 hours. The right plot summarizes all of the data with each dot representing a single well. a.u.: arbitrary units

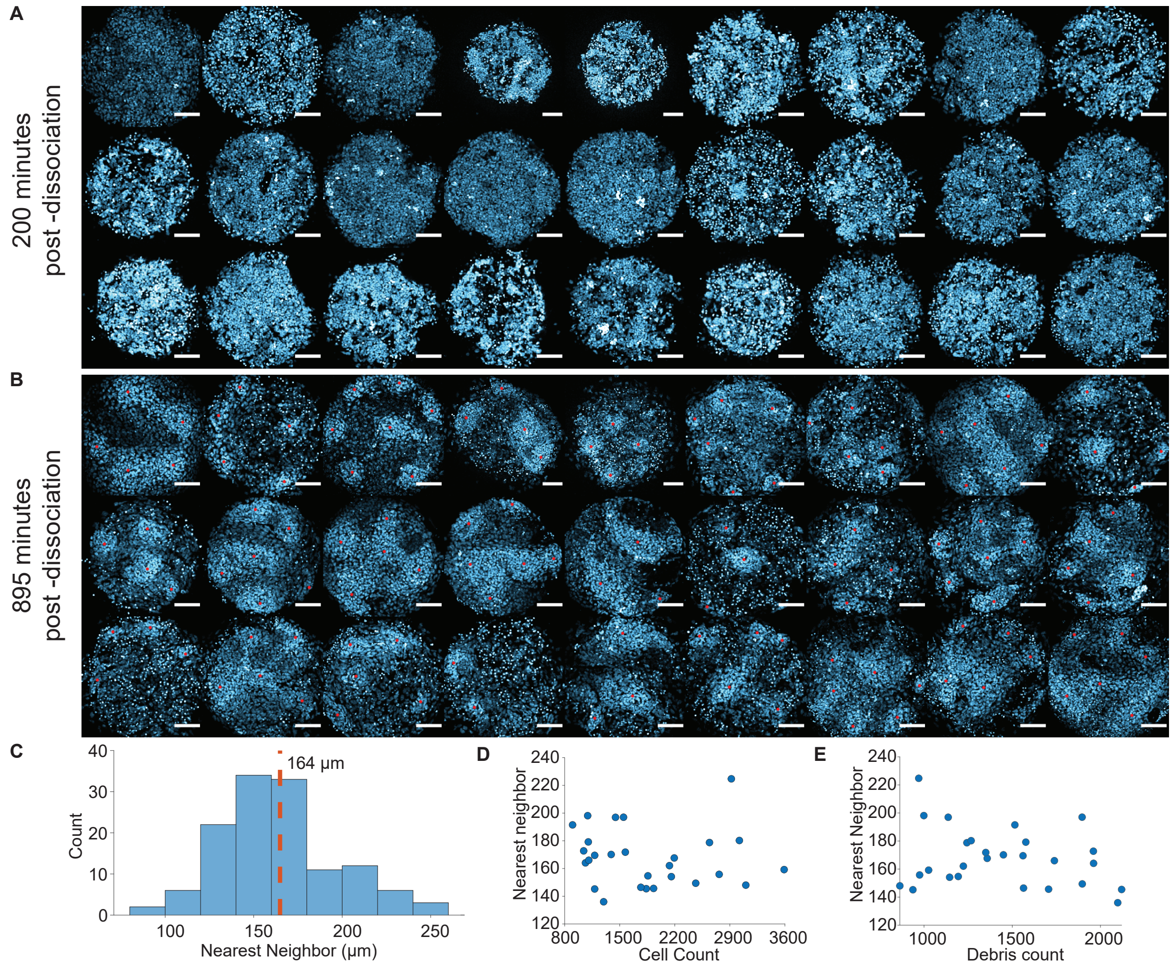


Figure S3: **Pattern appearance and spot spacing of Brachyury patterns in ePSMs. A)** Images of wells near the beginning of patterning in all 27 wells used in the density correlation analysis. **B)** $T$ pattern 700 minutes later of wells in A. Scale bars are 100 µm and intensity ranges are the same for the same wells in A and B. Red dots are manually selected points based on an apparent $T$ intensity peak. **C)** Nearest Neighbor Distribution of marked red spots in B. **D)** Relationship between average nearest neighbor distance and cell count across all wells. **E)** Relationship between average nearest neighbor distance and debris count across all wells.

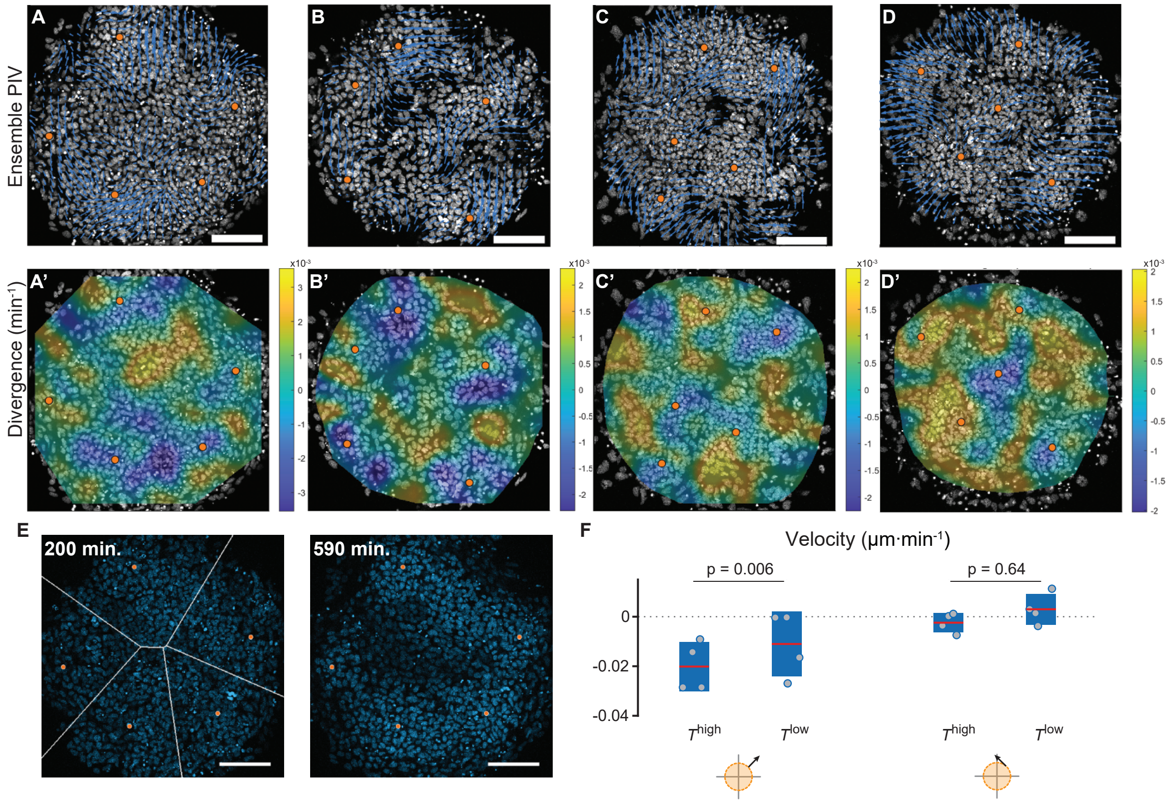


Figure S4: **Particle image velocimetry of tracked wells and association of aggregation-directed movement with *T* expression. A–D)** Four replicate wells used for cell tracking were subjected to ensemble particle image velocimetry (PIV) over a 200-minute interval from 395 to 595 minutes post-dissociation. Scale bar: 100 µm. **A'–D')** Divergence of the vector fields shown above. Manually selected aggregation points used for the radial velocity analysis in Fig. 1E are indicated by orange dots. **E)** To determine whether initial *T* reporter activity was associated with subsequent aggregation-directed movement, cells were assigned at 200 minutes to the nearest future centroid identified at 590 minutes. White lines mark the boundaries between the different 5 regions corresponding to cells associated with each of the 5 centroids. **F)** Cells from panel E were further divided into *T*-high and *T*-low populations according to whether their reporter activity at 200 minutes was above or below the median. Radial and tangential velocities were compared between the two populations. Points represent mean velocities from four replicate wells. Two-sided *p*-values were calculated for the differences between the T-high and T-low mean velocity components using spatial randomization procedure (*Methods*) in which the five centroids were randomly rotated and reflected relative to the cell tracks 2,000 times within each well.

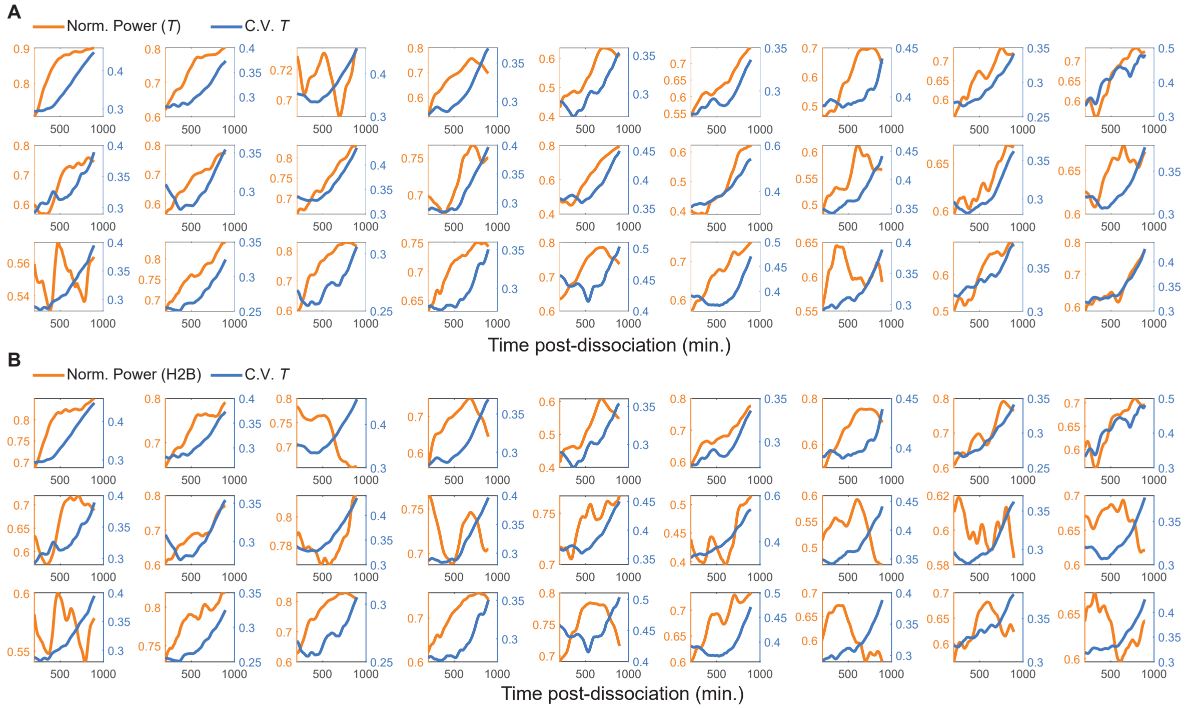


Figure S5: **Plots of individual wells comparing normalized power to coefficient of variation A.)** All 27 wells are shown with a comparison between normalized power of the $T$ reporter channel and C.V. of the Brachyury reporter distribution. **B.)** All 27 wells are shown with a comparison between normalized power of the H2B channel and C.V. of the Brachyury reporter distribution. All values plotted without standardization.

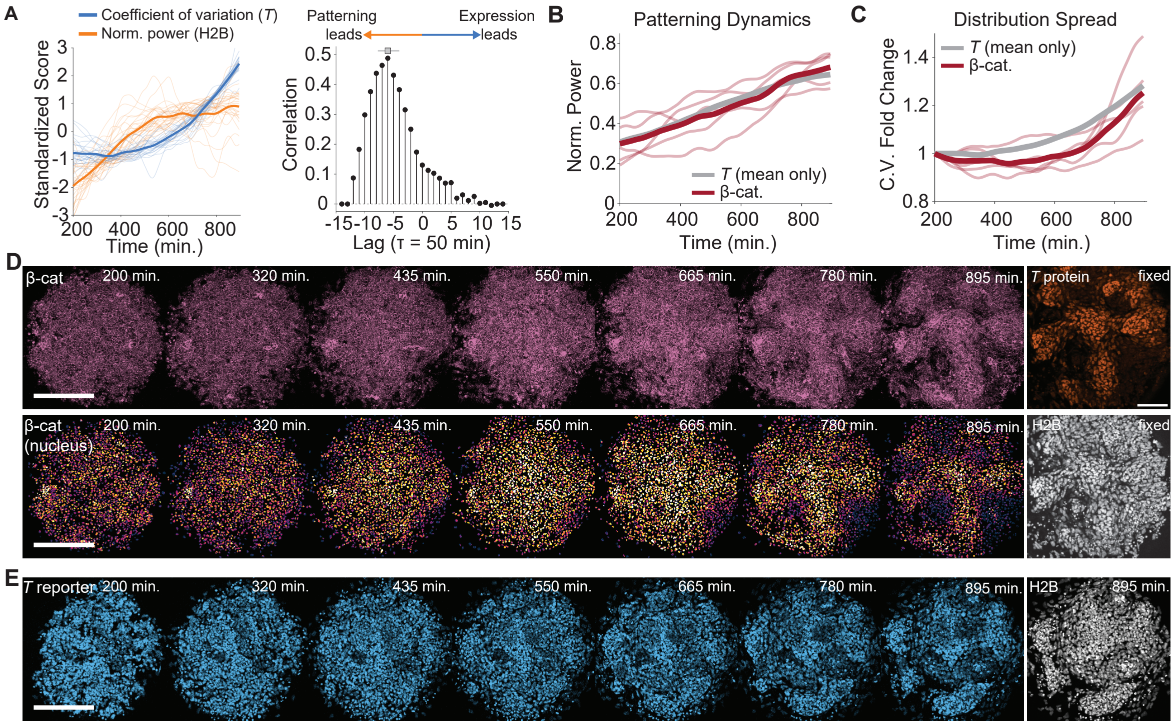


Figure S6: **$\beta$-catenin does not prepattern Brachyury domains A)** Comparison of standardized scores of the normalized power of cell nuclei patterns quantified from H2B reporter images (orange) to the $T$ distribution coefficient of variation (blue) for 27 wells. Cross-correlation analysis averaged across wells exhibited a peak lag at -300 minutes. (Gray square and line: median and inter-quartile range of cross-correlation peak positions of the individual wells.) **B)** Comparison of normalized power between $T$ and $\beta$-catenin images during ePSM formation. The mean $T$ curve comes from data presented in Fig. S5. No additional z-score standardization of the normalized power was applied in this comparison. **C)** Comparison of the fold change in the coefficient of variation of $T$ reporter activity and nuclear $\beta$-catenin intensity. Because CV values differed systematically between reporters, fold change was calculated independently for each well by dividing its coefficient-of-variation trace, without z-score standardization, by the value at the first analyzed time point. The mean $T$ trajectory was calculated from the same underlying CV traces presented in Fig. S5. **D)** Time lapse images of $\beta$-catenin patterning dynamics. Top row shows $\beta$-catenin images without subcellular emphasis while bottom row shows only the $\beta$-catenin nuclear pixel values obtained through nuclei segmentation. Scale bars are 200 µm. Endpoint immunostaining done on PFA fixed samples to confirm $T$ protein spot patterns. Cells were fixed after 980 minutes. Scale bar is 100 µm. **E)** Time lapse images of $T$ reporter patterning dynamics as a qualitative comparison to images in D. Scale bar is 200 µm.

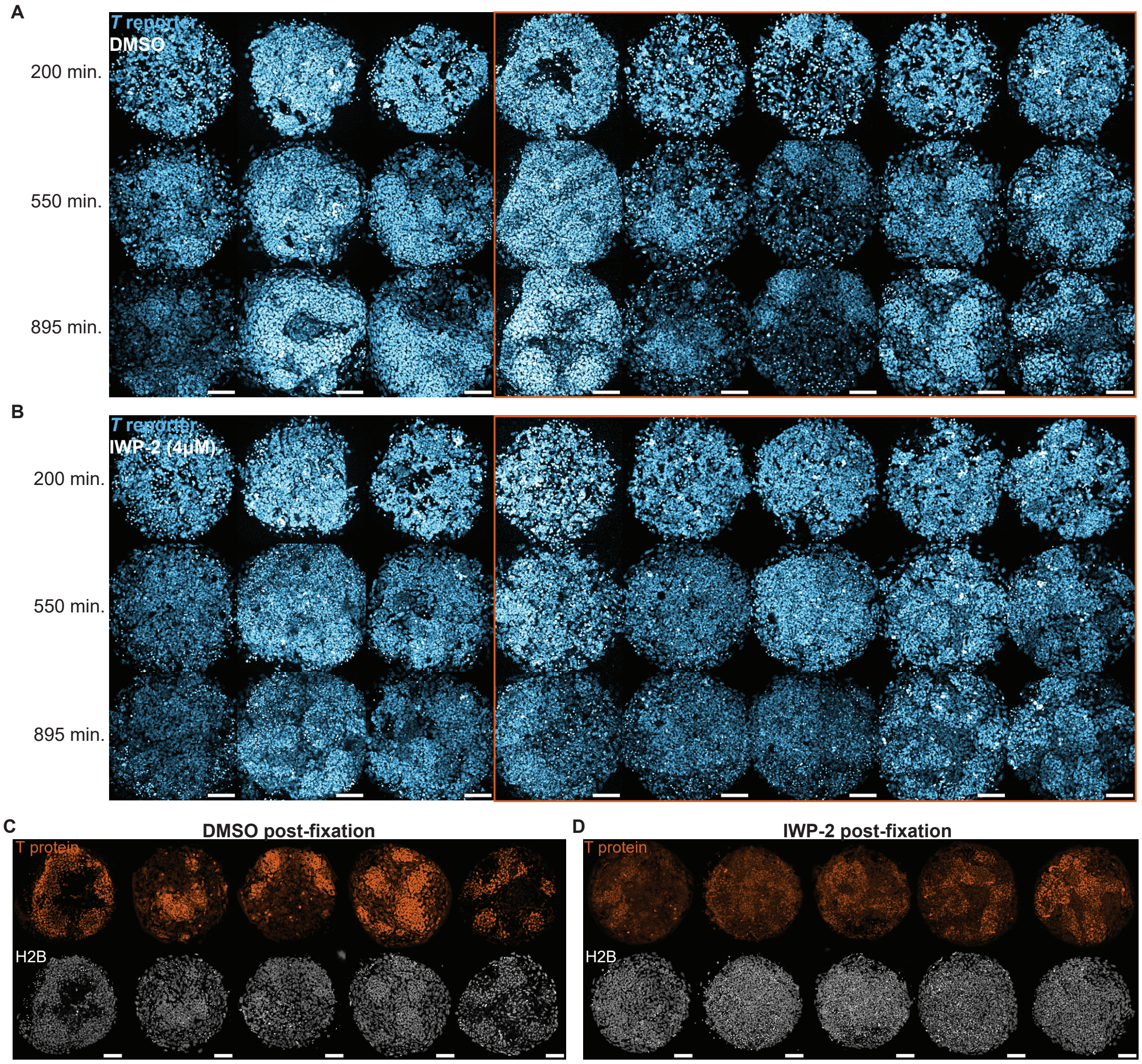


Figure S7: **Image examples of samples used in IWP-2 experiments A)** All 8 replicates for the control condition with labeled rows of beginning, middle, and end time points. Scale bars: 100 µm. **B)** All 8 replicates for the IWP-2 treatment condition with labeled rows of beginning, middle, and end time points. Scale bars: 100 µm. **C)** Immunostaining for Brachyury protein with corresponding images of cell nuclei (H2B-mcherry) for DMSO treated conditions. Fixed DMSO images correspond wells of the last 5 columns of panel A that are highlighted with an orange box and are most closely comparable to the images displayed in the 895 minute row. Fixation time from left to right: 1,145; 1,075; 1,075; 1,330; 1,330 minutes. **D)** Immunostaining for Brachyury protein with corresponding images of cell nuclei (H2B-mcherry) for 4 $\mu mol\,L^{-1}$ IWP-2 treated conditions. Fixed IWP-2 images correspond to wells of the last 5 columns of panel B that are highlighted with an orange box and are most closely comparable to the images displayed in the 895 minute row. Fixation time from left to right: 1,145; 1,075; 1,075; 1,330; 1,330 minutes. Scale bars: 100 µm.

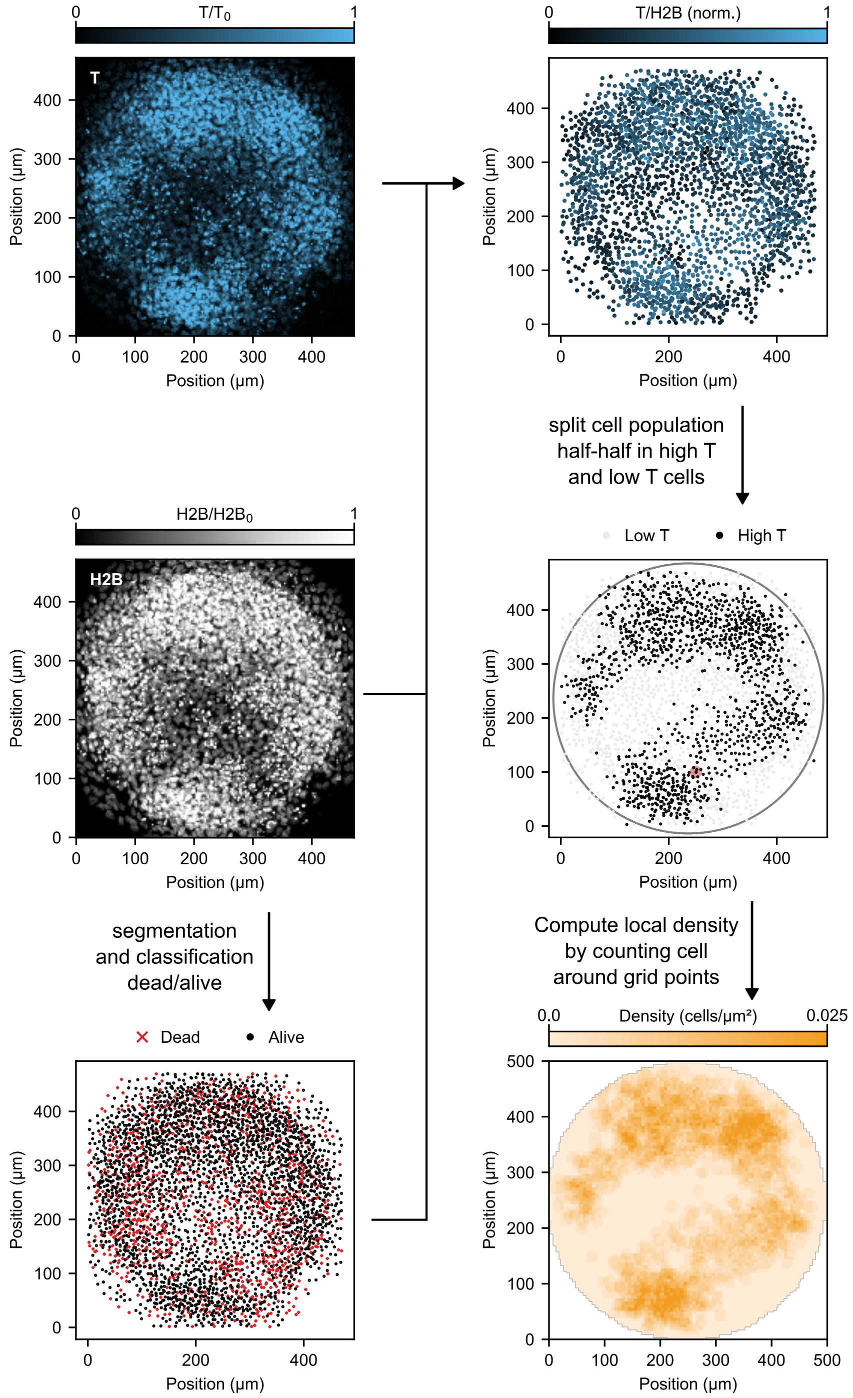


Figure S8: **Analysis pipeline to obtain the $T$+ density field.** $T$ gene activity is obtained by dividing the signal from the $T$ reporter by the signal in the H2B channel. Objects are further classified as dead or alive before thresholding by the median $T$ gene activity. Local density is then obtained by discretization of the image into a grid and counting the number of cells close to each grid point.

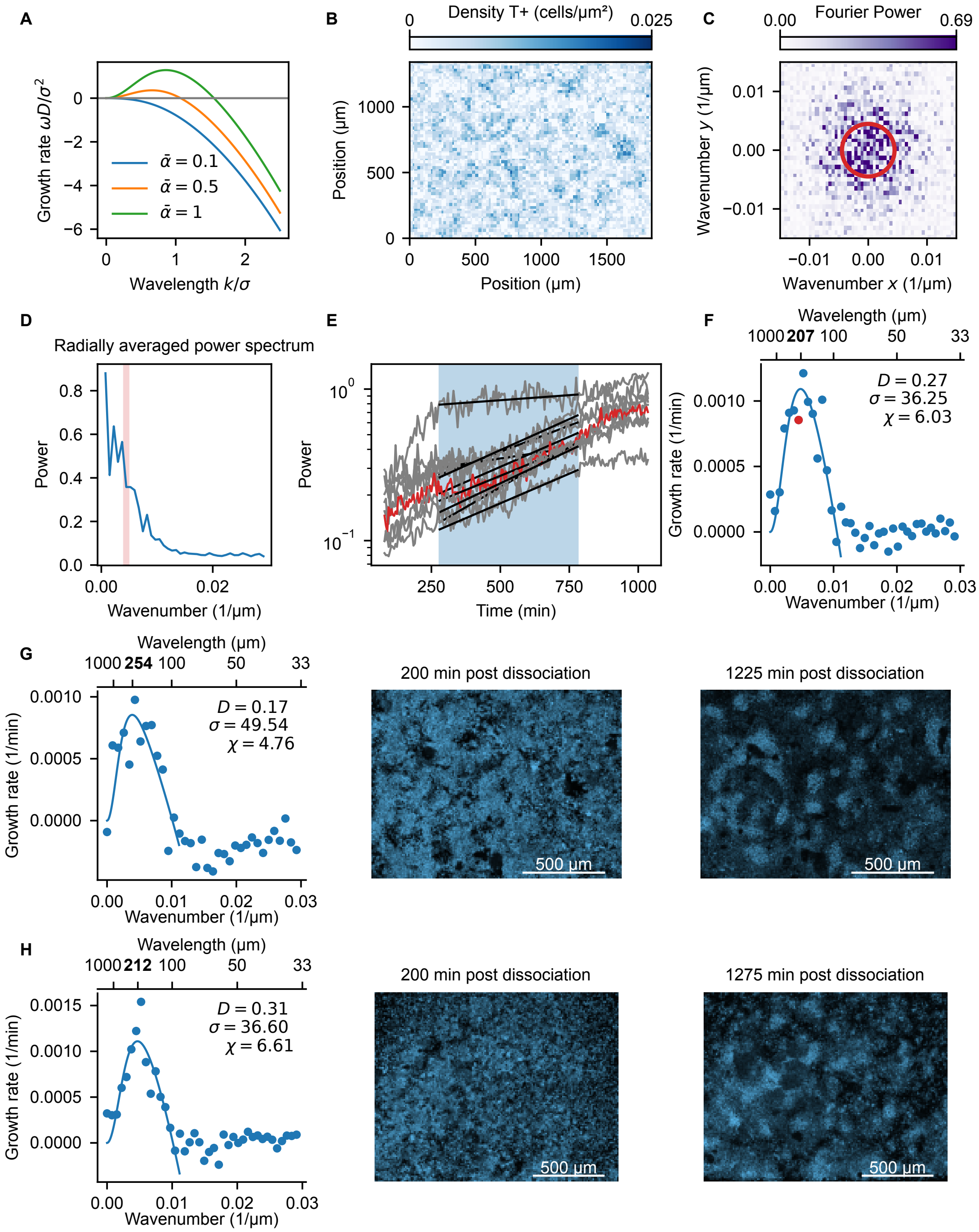


Figure S9: **Fitting the dispersion relationship A)** Theoretical dispersion relationship in scaled units, for different values of the normalized interaction strength $\bar{\alpha} = \rho_0(1 - \rho_0/\rho_{\max})(\chi\sigma)/D$. **B)** Example of a $T+$ density field obtained from one time point of a patterning experiment in large wells. **C)** The 2D Fourier spectrum of the image in B. Red circle correspond to all wave vectors with given wavenumber, corresponding to red highlights in panels DEF. **D)** The radially averaged power spectrum. **E)** Fitting an exponential function to power over time yields growth rates of each individual wavenumber. **F)** Dispersion relation: plot of growth rate as a function of wavenumber. The bold tick indicates the wavelength at which the fitted dispersion relation is maximal. **G-H)** The second and third replicates of the dispersion relationship fits.

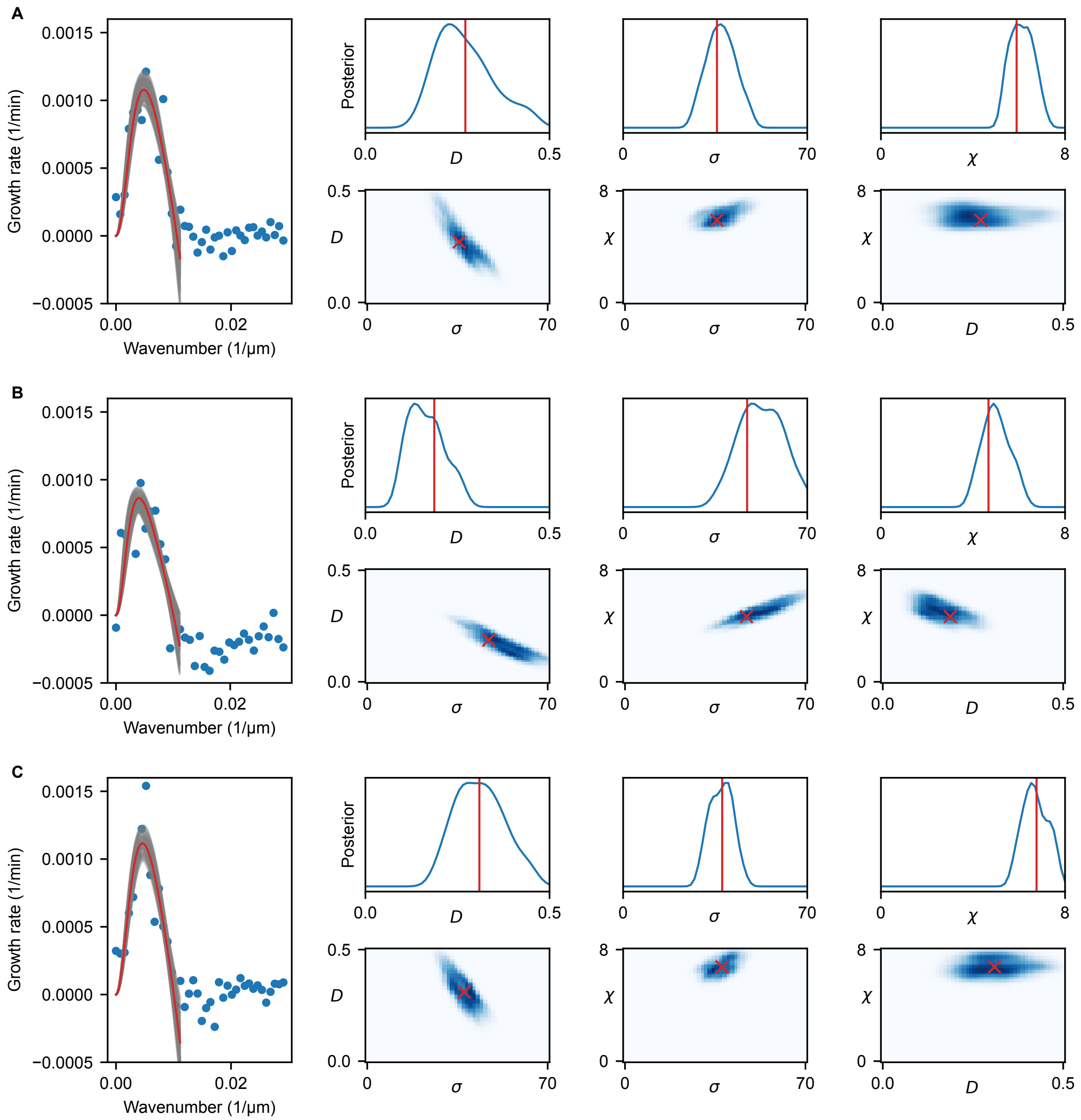


Figure S10: **Bayesian approach to fitting dispersion relationships A-C)** Three replicates with experimental dispersion relationship. Left: gray lines corresponding to the set of best-fit parameters resulting from the approximate Bayesian computation algorithm. Red line is the best fit. Right: top row shows the marginal posterior distribution of each parameter, bottom row shows the pairwise posterior distribution. Parameter units: $D$ ($\mu m^2\, min^{-1}$), $\chi$ ($\mu m^3\, min^{-1}\, cells^{-1}$), $\sigma$ ($\mu m$).

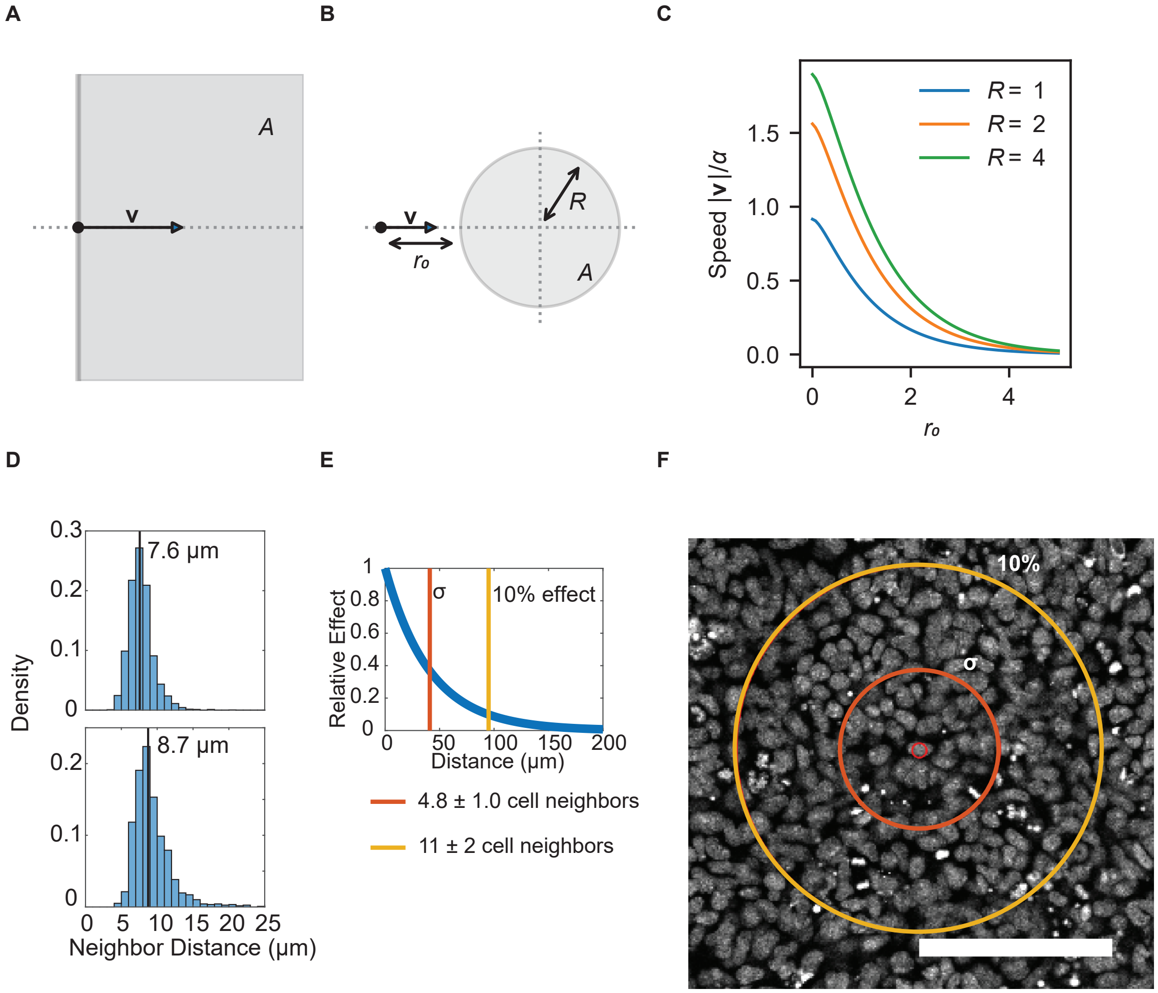


Figure S11: **Estimating the strength and range of cell-cell interactions A)** The maximal value of the velocity in the model is obtained with zero packing at a point next to a half plane filled with cells. In this case the integral that appears in the velocity calculation is equal to 2, and the speed in physical units is $2\rho\chi$ (with $\rho$ the constant density in the half plane). **B)** Diagram for the calculation of velocity at distance $r_0$ from a disk with radius $R$. **C)** Numerically computed values of the integral determining the velocity for the geometry in panel B. **D)** Histogram of nearest-neighbor distances between cells, two well-level examples shown. Black lines represent median of the distribution. Cell count for each example: 2500 cells (top) and 1,200 cells (bottom). The average and standard deviation of the nearest neighbor distribution was $8.5 \pm 0.7\,\mu\text{m}$ for $n = 4$ replicates. **E)** Relative interaction strength as a function of distance, with distance expressed in units of the mean nearest-neighbor spacing in the legend below. Uncertainties in the converted distances were calculated using standard error propagation for a ratio, assuming independent uncertainties in the interaction length and nearest-neighbor spacing. **F)** Circles with radii corresponding to panel E overlaid on top of an image of H2B-mCherry expressing cells to illustrate the effective interaction distances. Small red circle marks the centroid of the cell that acts as the center for the comparison. Scale bar = $100\,\mu\text{m}$.

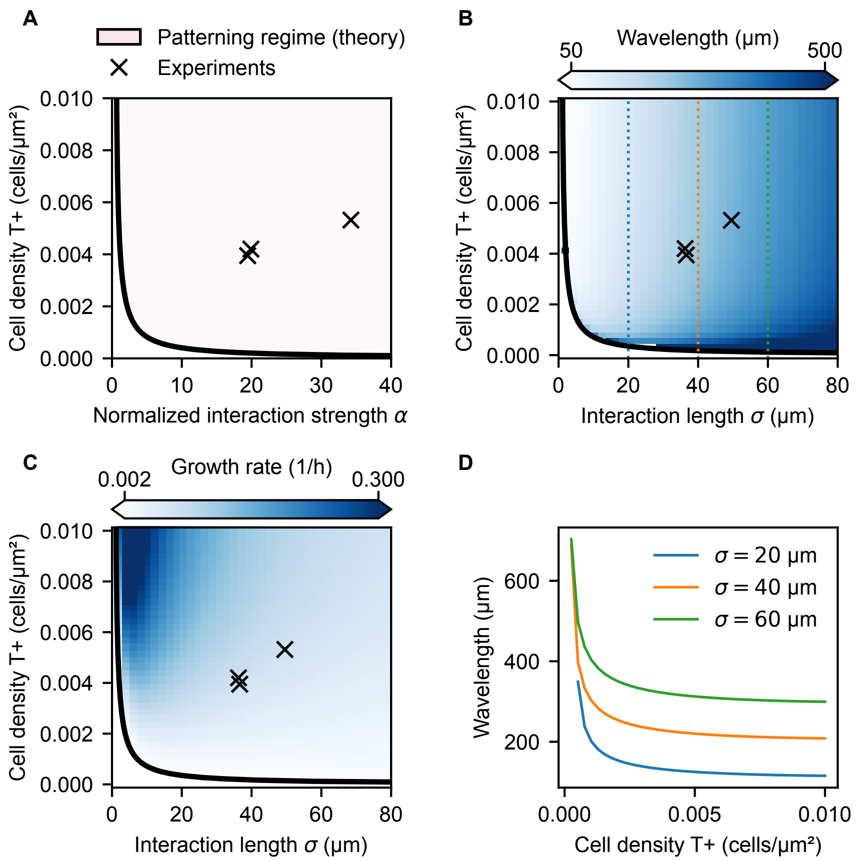


Figure S12: **Linear stability analysis of nonlocal continuum model gives patterning threshold and estimates of initial space and time scales A)** Patterning regime resulting from the linear stability analysis. Patterns are possible if the normalized interaction strength $\alpha$ and cell density are sufficiently large. Fitted parameter values corresponding to experiments are indicated with crosses. **B)** The dominant wavelength of the initial pattern as a function of cell density and interaction length. **C)** Growth rate of dominant spatial mode as function of cell density and interaction length during initial patterning. In B and C, wavelength and growth rate are calculated using the average values of $D$ and $\chi$ from the dispersion relation of the experimental data. Panels A–C: vertical axis is the spatially homogeneous mean T+ density $\rho_0$ and the crosses indicate values estimated from the three large-well experiments shown in Fig. 3B and Fig. S9. **D)** Wavelength as a function of density for selected values of interaction length scale $\sigma$, corresponding to the vertical dashed lines in panel B.

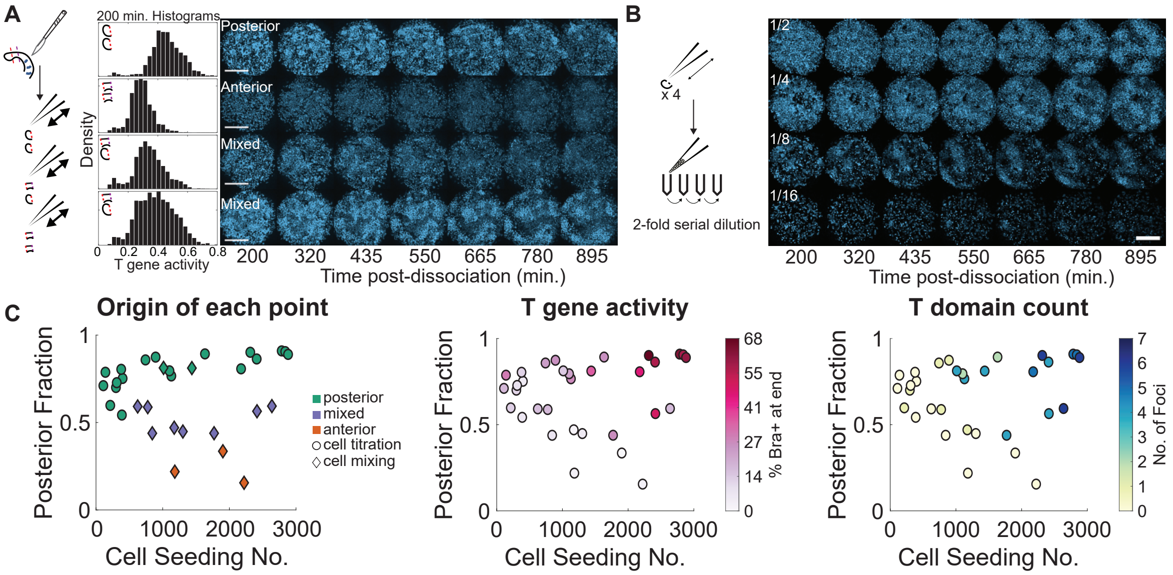


Figure S13: **Influence of initial conditions on patterning A)** Cell mixing experiments were conducted by mixing together different fractions of posterior and anterior cells obtained from the mouse embryonic tail. Schematic is presented alongside the histograms of $T$ reporter distribution at the start of the experiment. Images show the time evolution of the patterns for each condition. Scale bars are 100 µm **B)** Cell titration experiments where the tailbud/posterior PSM fraction is serially diluted before seeding into wells. Schematic describing the experiment is presented alongside one experimental replicate. Scale bar is 100 µm **C)** The combined data from cell mixing and cell titration experiments to generate an initial conditions regime diagram. $T$ gene activity is measured by taking the fraction of cells above a given $T$ gene activity threshold at the end of the experiment. Brachyury domain count is obtained by manual counting of spots of high Brachyury gene activity.

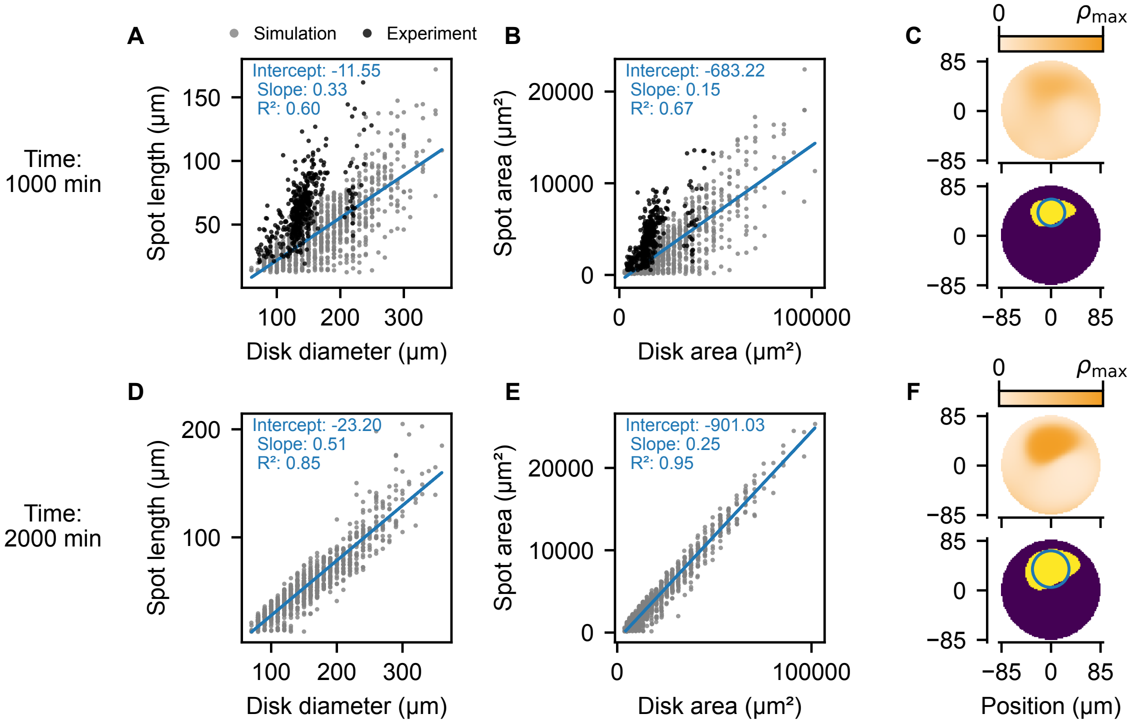


Figure S14: **Scaling of spot size as a function of system size A)** Linear dimensions of single spot as a function of system size at 800 min of simulation time. The experimental dataset also shown in Fig. S1 is added on top. The line and inset text correspond to the linear fit to the simulation points. **B)** Area of single spot as a function of system size at a timepoint comparable to 1000 min after cell seeding (in particular, 800 min of simulation time, with initial conditions based on the experimental density distribution at 200 min post seeding, see also Methods). **C)** We computed the spot size by segmenting the simulation output (threshold: $\rho_{\max}/2$). We performed 100 simulations per disk size and for this analysis kept only simulations that showed a single spot, which means for larger systems there are fewer points since we mainly observed multispot patterns for those larger systems. The length of a spot was calculated as the minimum Feret diameter. **D,E,F)** Same as panels A, B, C but at a later timepoint (2000 minutes).

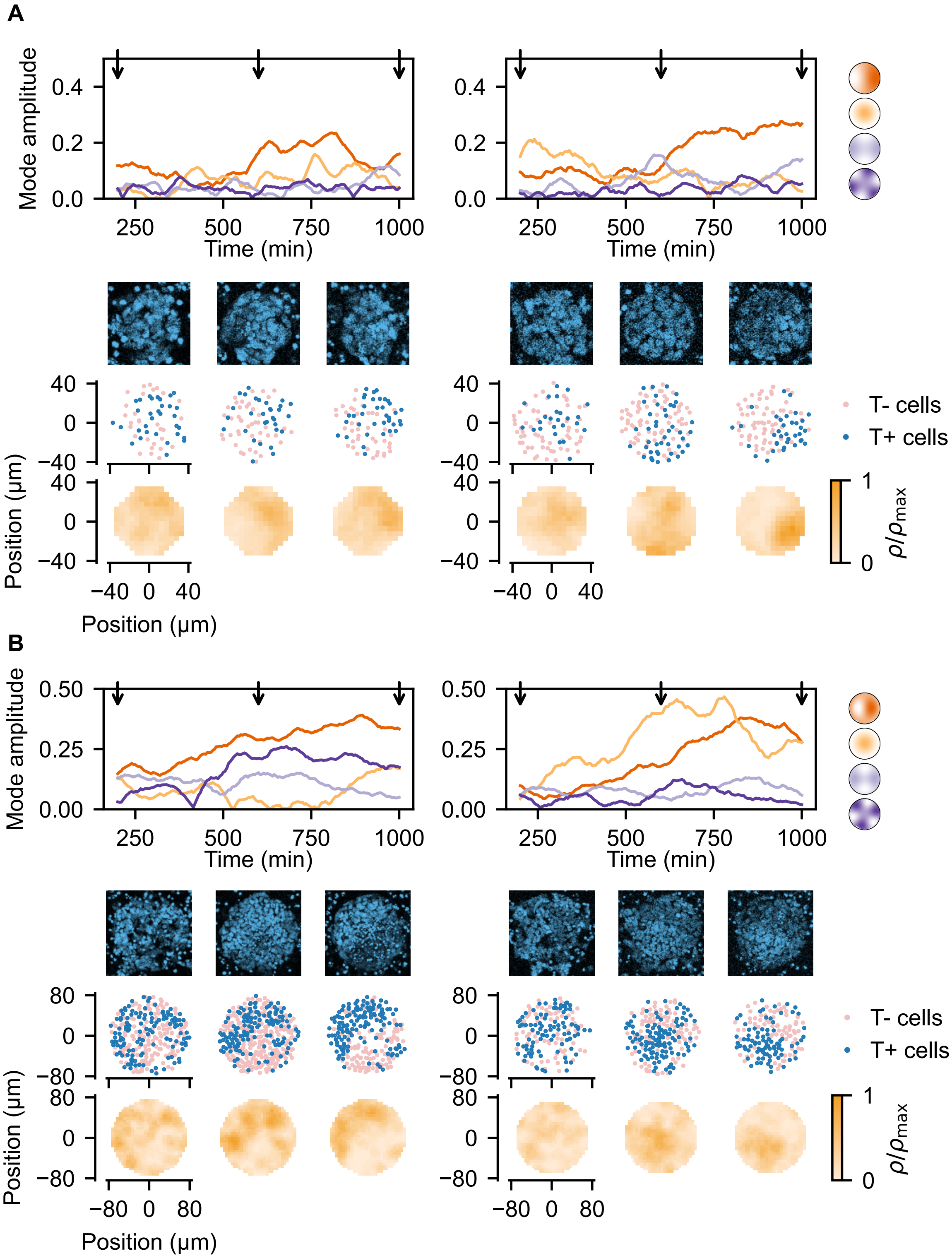


Figure S15: **Examples of mode evolution on selected disks A)** Two representative 80 μm diameter disks. **B)** Two representative 160 μm diameter disks. Top panel shows the amplitudes of the four indicated modes over time, arrows indicate the snapshot times shown below. Snapshots of (top to bottom) the $T$ reporter intensity, the positions of high and low $T$ cells after segmentation and classification, and the calculated density of $T+$ cells.

### Movie Legends

**Movie S1. Dynamics of *T* patterning.** Time-lapse video of pattern formation in the *T* reporter channel beginning 200 minutes post tissue dissociation. Example z-slice is 5/13. Images captured at 5-minute intervals and scale bar = 100 µm.

**Movie S2. Before JLY washout.** Composite time-lapse video of *T* reporter (blue) overlaid on top of H2B-mCherry (gray) to show arrest of cell movement. Imaging begins 51 minutes after tissue dissociation and the addition of JLY. Washout is performed immediately after the last imaged frame. Video corresponds to the example in Fig. 1H from z-slice 3/13. Images captured at 5-minute intervals and scale bar = 100 µm.

**Movie S3. After JLY washout.** Composite time-lapse video of *T* reporter (blue) overlaid on top of H2B-mCherry (gray) to show recovery of cell movement. Imaging begins shortly after washout of JLY and ends 1,000 minutes after tissue dissociation. Video corresponds to the example in Fig. 1H from z-slice 3/13. Images captured at 5-minute intervals and scale bar = 100 µm.

**Movie S4. Dynamics of *T* patterning in large wells.** Top half of the video depicts pattern formation in the *T* reporter channel in large wells. In this example, cells are pooled together from 8 embryos. Bottom half corresponds to the *T*+cell density field after processing. Images tiled in a non-overlapping manner, captured at 5-minute intervals and scale bar = 100 µm.

**Movie S5. Dynamics of *T* patterning on small disks.** Composite time-lapse video of *T* reporter (blue) overlaid on top of H2B-mCherry (gray) to show examples of cell confinement on micropatterned disk substrates. In the video, three examples of 80 µm disks are shown in the top row and three examples of 160 µm disks are shown in the bottom row. Videos were captured at 5-minute intervals and scale bar = 50 µm.

**Movie S6. Dynamics of *T* patterning at longer times.** Time-lapse videos of pattern formation in the *T* reporter channel beginning 650 minutes post tissue dissociation and extending to 1,250 minutes. The left example is an extended version of Movie S1 while the right example is a separate example from another experimental day. White arrows are added annotations marking merging events. Example z-slice is 5/13. Images captured at 5-minute intervals and scale bar = 100 µm.
